# Anti-Aβ immunotherapy-mediated amyloid clearance attenuates microglial activation without inducing exhaustion at residual plaques

**DOI:** 10.1101/2025.04.10.645950

**Authors:** Lis de Weerd, Selina Hummel, Stephan A. Müller, Iñaki Paris, Thomas Sandmann, Marie Eichholtz, Robin Gröger, Amelie Englert, Stephan Wagner, Connie Ha, Sonnet S. Davis, Valerie Warkins, Dan Xia, Brigitte Nuscher, Anna Berghofer, Marvin Reich, Astrid Feiten, Kai Schlepckow, Michael Willem, Stefan F. Lichtenthaler, Joseph W. Lewcock, Kathryn M. Monroe, Matthias Brendel, Christian Haass

## Abstract

Anti-amyloid β-peptide (Aβ) immunotherapy was developed to reduce amyloid plaque pathology and slow cognitive decline during progression of Alzheimer’s disease. Efficient amyloid plaque clearance has been proven in clinical trials testing anti-Aβ antibodies, with the impact on cognitive endpoints correlating with the extent of plaque removal. However, treatment is associated with adverse side-effects, such as oedema and haemorrhages, which are potentially linked to the induced immune response. To improve the safety profile of these molecules, it is imperative to understand the consequences of anti-Aβ antibody treatment on immune cell function. Here, we investigated the effects of long-term chronic anti-Aβ treatment on amyloid plaque pathology and microglial response in the APP-SAA triple knock-in mouse model. Mice were treated weekly with anti-Aβ antibody from 4-8 months of age. Long-term treatment with anti-Aβ results in a robust and dose-dependent removal of amyloid plaque pathology, with a higher efficiency for removing diffuse over dense-core plaques. Analysis of the CSF proteome indicates a reduction of markers for neurodegeneration including Tau and α-Synuclein, as well as immune cell related proteins. Bulk RNA-seq revealed a dose-dependent decrease in brain-wide disease-associated microglial (DAM) and glycolytic gene expression, which is supported by a parallel decrease of glucose uptake and protein levels of Triggering receptor of myeloid cells 2 (Trem2) protein, a major immune receptor involved in DAM activation of microglia. In contrast, DAM activation around remaining plaques remains high regardless of treatment dose. In addition, microglia surrounding remaining plaques display a dose-dependent increase in microglial clustering and a selective increase in antigen presenting and immune signalling proteins. These findings demonstrate that long-term chronic anti-Aβ mediated removal of Aβ leads to a dose dependent decrease in brain-wide microglial DAM activation and neurodegeneration, while microglia at residual plaques display a combined DAM and antigen presenting phenotype that suggests a continued treatment response.

Graphical abstract: Schematic overview of the effects of chronic long-term anti-Aβ treatment in APP-SAA mice Schematic was created with BioRender.com

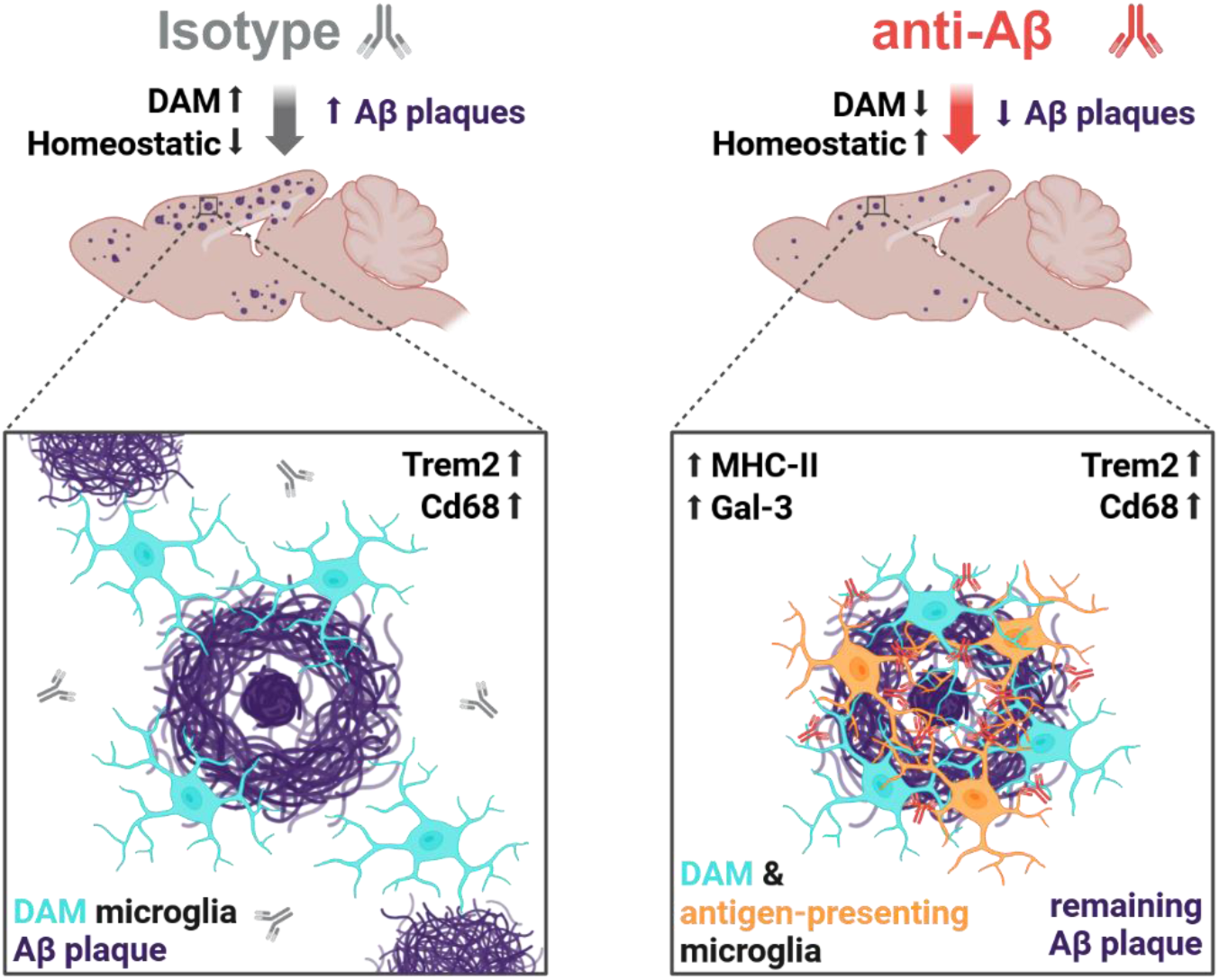

## Introduction

Alzheimer’s disease (AD) is characterized by the deposition of aggregation-prone amyloid-β-peptide (Aβ). In patients suffering from autosomal dominant AD, it has been shown that aggregation of Aβ is initiated approximately 25 years before symptom onset, triggering a pathological cascade of Tau aggregation, neurodegeneration and neuroinflammation that ultimately leads to cognitive decline ^1,2^. Based on the pioneering work by Schenk and colleagues ^3^, therapeutic antibodies, such as Aducanumab, Lecanemab and Donanemab were developed that target aggregated forms of Aβ. In phase III clinical trials, these antibodies have been shown to significantly reduce amyloid plaque burden upon chronic treatment in patients and, for the latter two to modestly slow cognitive decline ^4–6^. In addition, pathological Tau is reduced and patients with low Tau burden show stronger clinical effects, supporting the idea that early amyloid removal can alter the course of the pathological cascade. Anti-Aβ immunotherapy is the first and currently only disease-modifying treatment for patients with early symptoms of AD or mild cognitive impairment (MCI). However, beyond established clinical outcomes, the effects of early and long-term immunotherapy remain largely unknown.

Aβ-targeting antibodies remove Aβ by recruiting the brain’s immune system, namely microglia, inducing a transient immune response that results in the phagocytosis of aggregated Aβ ^7,8^. This treatment appears to bypass many of the risks associated with active immunization ^9^, but is associated with side-effects such as brain oedema and haemorrhages (detected with MRI as amyloid related imaging abnormalities (ARIA)), which can have fatal consequences in rare cases ^10^. The mechanisms of these side-effects are currently poorly understood, but are potentially linked to immune cell activation^11,12^. To improve the safety of anti-Aβ antibody treatment, it is therefore important to gain a deeper understanding of the long-term effects of anti-Aβ antibody treatment on microglia.

Microglia can exist in transient states that are defined by transcriptional signatures, which signify their functions. Under pathological conditions, such as the accumulation of aggregated Aβ, microglia transition from a homeostatic state into a disease-associated microglia (DAM) state that is associated with increased metabolic activity, proliferation, chemotaxis and phagocytosis ^13,14^. This transition relies in part on the presence of microglial Triggering Receptor in Myeloid Cells 2 (TREM2). Loss of function variants of TREM2 are associated with an increased risk for late onset AD (LOAD)^15,16^ and microglia that lack TREM2 do not convert into the full DAM state and are less capable to deal with Aβ-related challenges ^17–22^. Conversely, higher CSF TREM2 is associated with slower rates of Aβ deposition, reduced cortical shrinkage and diminished cognitive decline ^23–26^. The microglial TREM2-dependent transition into the DAM state is therefore thought to be a protective response to fight Aβ deposition and is currently explored for the development of TREM2 agonists ^13^.

Lack of Trem2 was previously shown to impact the efficacy of anti-Aβ antibody treatment *ex vivo* ^27^, suggesting that microglial function can influence the efficacy of treatment. Vice versa, Aβ-targeting immunotherapies can affect microglia, which has already been considered in early immunotherapy studies to a limited extent ^7,8,28^. In recent years, studies have shown that short-term treatment (<4 weeks) with high doses of anti-Aβ antibody in mouse models of amyloidosis results in little to no reduction of Aβ load, but is associated with increased expression of homeostatic genes and reduced expression of DAM genes ^29,30^. Studies on long term-treatment with anti-Aβ (>8 weeks) also report limited and variable effects on Aβ removal in a dose-dependent manner, which is associated with increased clustering of microglia around plaques ^30–34^. However, these studies provide little insights into the long-term treatment effects on the transcriptional and functional state of microglia or potential biomarker readouts for monitoring treatment response.

Here, we investigate the effect of early chronic anti-Aβ antibody (referred to as anti-Aβ) treatment, on Aβ deposition and microglial activation in APP-SAA knock-in (KI) mice ^35^. After 16 weeks of treatment, anti-Aβ reduced plaque load, with a higher efficiency for loosely aggregated fibrils, and concomitantly reduced neuritic dystrophies in a dose-dependent manner. This was accompanied by a reduction in biomarkers of neuritic dystrophy and microglial activation in CSF. Bulk-RNA sequencing, in vivo PET-imaging together with proteomic analyses demonstrates a dose-dependent global attenuation of DAM activation and glycolysis, but identifies increased expression of genes associated with antigen presentation. When analysing microglia at residual plaques, we find increased clustering, unchanged DAM activation, but the induction of MHC-II and Galectin-3 in a dose-dependent manner.

## Methods

### Mice

All animal experiments were approved by the Ethical Review Board of the Government of Upper Bavaria. Mice were group housed with littermates on a normal 12-hour light/dark cycle with ad libitum access to food and water. Both genders were used for all experiments. APP-SAA^ki/ki^ x hTfR^ki/ki^ (APP-SAA x hTfR KI) mice^35–37^ were acquired from Denali Therapeutics or bred in our mouse facility and maintained on a C57BL/6J genetic background. hTfR KI was bred into these mice in preparation of future antibody dosing studies that exploit antibody transport vehicle (ATV) technology, but was not investigated in the current study and was previously not found to impact microglia phenotypes in response to Aβ ^37^ (and Fig. S1). Shipped mice were acclimated for a minimum of two weeks before entering experiments. All mice were housed in standard sized individually ventilated cages, with enriched environment and ad libitum access to food and water. Mice were maintained on a 12-hour light/dark cycle. For anti-Aβ treatment, the chimeric anti-amyloid antibody Aducanumab was used, which contains a mouse IgG2 Fc domain with full effector function ^31^. For isotype control, antibody 4D5 was used which has a mouse IgG2 Fc domain and is raised against human HER2,a non-existing target in mice ^38^. Mice were randomly assigned to a treatment arm: isotype antibody 1 mg/kg for cohort 1 (used for FBB-PET, immunofluorescent and protein analyses) and 10 mg/kg for cohort 2 (used for microglial RNA-seq and lipidomics, FDG-PET, CSF proteomic analyses), anti-Aβ 1 mg/kg, anti-Aβ 3 mg/kg or anti-Aβ 10 mg/kg. Mice were treated from the average age of 4.5 months by weekly intraperitoneal (i.p.) injection of antibody, which was thawed at 4°C and diluted with phosphate-buffered saline (PBS). Mice were sacrificed 7 days after the last antibody injection at an average age of 8.5 months.

### Small animal PET/MRI

All rodent PET procedures followed an established standardized protocol for radiochemistry, acquisition times and post-processing^39^, which was transferred to a novel PET/MRI system^40^. In brief, [^18^F]-FBB-PET (florbetaben) and [^18^F]-FDG-PET (fluorodeoxyglucose) were used to measure fibrillar amyloidosis and glucose metabolism respectively after antibody treatment. We studied PET images of 8,2 ± 0,2 month old APP-SAA mice (n=32) for FBB-PET and 8,5 ± 0,5 month old APP-SAA mice (n=35) for FDG-PET using at least n=8 per treatment group and tracer. All mice were scanned with a 3T Mediso nanoScan PET/MR scanner (Mediso Ltd, Hungary) with a triple-mouse imaging chamber. Isoflurane anaesthesia was applied for all PET experiments (1,5% at time of tracer injection and during imaging; delivery 3,0 L/min). Two 2-minute anatomical T1 MR scans (sagittal and axial) were performed after tracer injection (head receive coil, matrix size 96 × 96 × 22, voxel size 0,24 × 0,24 × 0,80 mm³, repetition time 677 ms, echo time 28,56 ms, flip angle 90°). Injected dose was 13,1 ± 2,1 MBq for [^18^F]-FBB and 19,1 ± 1,5 MBq for [^18^F]-FDG delivered in 200 µl saline via venous injection. PET emission was recorded in a dynamic 0-60 min window for FBB-PET and in a static 30-60 min window for FDG-PET. List-mode data within 400-600 keV energy window were reconstructed using a 3D iterative algorithm (Tera-Tomo 3D, Mediso Ltd, Hungary) with the following parameters: matrix size 55 × 62 × 187 mm³, voxel size 0,3 × 0,3 × 0,3 mm³, 8 iterations, 6 subsets. Decay, random, and attenuation correction were applied. The T1 image was used to create a body-air material map for the attenuation correction. Framing for FBB-PET was 6×10s, 6×30s, 6×60s, 10×300s.

All analyses were performed by using PMOD software (version 3.5, PMOD Technologies, Basel, Switzerland). To normalize FBB-PET data we generated V_T_ images with an image derived input function ^41^ ^42^, using the methodology described by Logan *et al.* implemented in PMOD^43^. The plasma curve was obtained from a standardized voxel of interest (VOI) placed in the myocardial ventricle. A maximum error of 10% and a V_T_ threshold of 0 were selected for modelling of the full dynamic imaging data. Normalization of the injected activity for FDG-PET was performed by generating standardized uptake values (SUV), reflecting the common read-out in clinical setting. A cortical volume-of-interest (comprising 40,9 mm³) was designed and served for extraction of FBB-PET values. FDG-PET values were extracted from a bilateral entorhinal VOI (compromising 13,0 mm³) which was delineated by regions of the Mirrione atlas^44^.

### Mouse brain, CSF and blood sampling

CSF collection was performed as described previously ^45^. Briefly, mice were anesthetized using a mix of medetomidine [0.5 mg/kg], midazolam [5 mg/kg], and fentanyl [0.05 mg/kg] (MMF). After complete anaesthesia, mice were head-fixed in a stereotax and the cisterna magna was surgically exposed. The dura was punctured using a borosilicate glass capillary (Sutter, B100-75-10) attached to medical grade tubing and CSF was gently extracted by applying backpressure on the tubing using a syringe. CSF samples were deposited from the capillary into protein Lo-Bind tubes (Eppendorf, 0030108094) and kept on ice until centrifugation at 2000g for 10 min at 4°C to pellet any red blood cells to control for contamination. After CSF collection, blood was extracted via cardiac puncture using a syringe, inserted into Microvette® 500 EDTA K3E tubes (Sarstedt, 20.1341.100), slowly inverted 10 times and kept on ice. Within 1 h, blood was centrifuged at 12700 rpm at 4°C for 10 minutes. Plasma was then transferred to a protein Lo-Bind tube and snap frozen. Mice were perfused via cardiac puncture with ice-cold PBS. Brains were split into two hemispheres and one hemisphere was fixed in 4% paraformaldehyde (PFA) for 48 hours. The other hemisphere was snap-frozen in liquid nitrogen and stored at -80°C.

### Immunofluorescence staining of mouse brain

50 μm brain sections were cut using a vibratome and stored in 15% glycerol + 15% ethylene glycol in PBS for 2 days at 4°C, before transferring to -20°C freezer for long-term storage. For immunostaining, free-floating sections were washed 5x in PBS on a shaker to remove storage medium. Antigen retrieval was performed in citrate buffer (pH6) or Tris-EDTA buffer (pH8 or pH9) at 80°C or 95°C for 30 min, depending on the antibody. After antigen retrieval, sections were cooled down to room temperature (RT), briefly washed in PBS and incubated in 10% normal donkey serum (NDS) in PBS + 0.3% Triton-X (blocking solution) on a shaker for 1-1.5 h.

Section were incubated overnight in blocking solution containing primary antibodies. The next day, sections were washed 3x in PBS + 0.3% Triton-X and incubated in secondary antibodies in blocking solution for 1-2 h. In case of co-staining with Thiazine Red (Morphisto, 12990.001) dye was added to the secondary solution, sections were washed 3x in PBS + 0.3% Triton-X. For Methoxy-X04 (MX-04, Tocris, 4920) co-staining, sections were incubated in 50% EtOH in PBS with MX-04 for 30 min at RT, washed 5 min in 50% EtOH in PBS and washed 3x in PBS. In case of HS169 (courtesy of Peter Nilsson, Linsköping University, Sweden) staining, dye was incubated 1:2500 in PBS for 15 min and washed 3x in PBS. If applicable, 40,6-diamidino-2-phenylindole (DAPI) was added to secondary antibody solution (1:1000). Sections were mounted onto Superfrost Plus slides with ProLongTM Gold antifade reagent (Thermo Fisher, P36980) or Fluoromount-G™ (Thermo Fischer, 00-4958-02). After 24 hours of drying, slides were stored at 4°C.

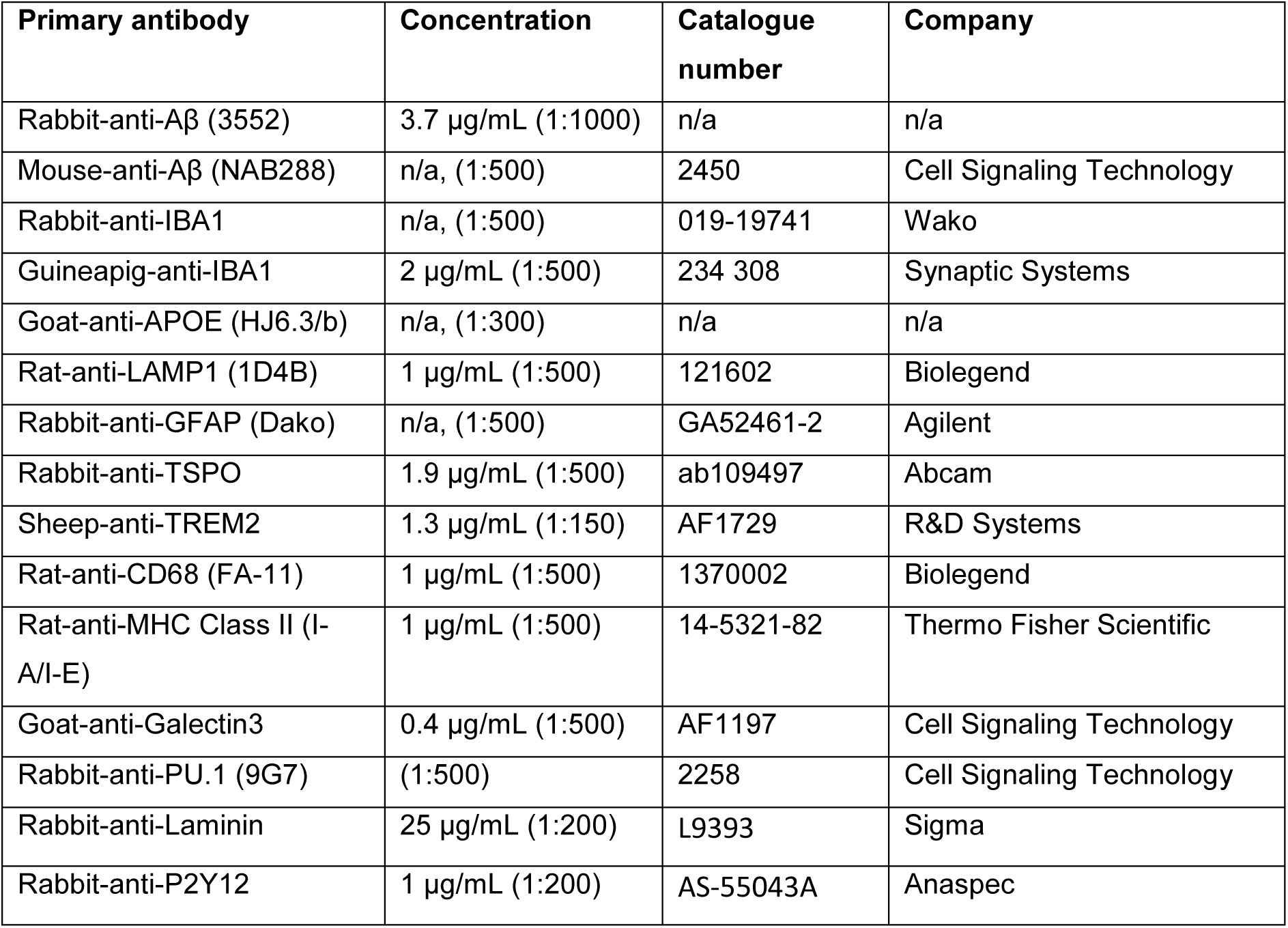

| Secondary antibody | Concentration | Catalogue number | Company |
| --- | --- | --- | --- |
| Donkey-anti-rabbit Alexa Fluor™ Plus 488 IgG (H+L) | (1:1000) | A32790 | Invitrogen |
| Donkey-anti-rabbit Alexa Fluor™ Plus 647 IgG (H+L) | (1:1000) | A32790 | Invitrogen |
| Donkey-anti-mouse Alexa Fluor™ Plus 488 IgG (H+L) | (1:1000) | A32766 | Invitrogen |
| Donkey-anti-mouse Alexa Fluor™ Plus 647 IgG (H+L) | (1:1000) | A32787 | Invitrogen |
| Donkey-anti-rat Alexa Fluor™ Plus 647 IgG (H+L) | (1:1000) | A32795 | Invitrogen |
| Donkey-anti-goat Alexa Fluor™ Plus 647 IgG (H+L) | (1:1000) | A32849 | Invitrogen |
| Donkey-anti-sheep Alexa Fluor™ Plus 647 IgG (H+L) | (1:1000) | A21448 | Invitrogen |
| Donkey-anti-guinea pig Alexa Fluor™ Plus 647 IgG (H+L) | (1:1000) | A21450 | Invitrogen |

### Prussian Blue staining of hemosiderin deposits in mouse brain

For quantification of hemosiderin deposits, slides were mounted onto Superfrost Plus slides and dried for 2 h at room temperature (RT). Slides were rehydrated in PBS and incubated in Prussian blue solution (2g potassium hexacyanoferrate (II) trihydrate (Sigma, P9387) in 100 mL dH_2_O) for 20 min and in 0.1% Nuclear Fast Red solution (Morphisto, 10264.00500) for 5 min and washed in dH_2_O. Slides were dehydrated from 70% to 100% EtOH and mounted using VectaMount® Express Mounting Medium (Vector Labs, VEC-H-5700). The number of Prussian Blue deposits was quantified from 5 brain sections of each mouse by stereology using the Leica DMi8 fluorescence microscope. Images of deposits were acquired using a 40x air lens (0.65NA, Leica). Area and number of observed deposits was quantified from images using ImageJ ^46^.

### Microscopy and image acquisition

Epifluorescence images were acquired with a Leica DMi8 equipped with a mercury lamp (EL6000, Leica) using a 20x air lens (0.4NA, Leica) or an Olympus VS200 Slideview slide scanner using a 20x air lens (0.8NA, 0.274 μm/pixel). Leica scanned tiles were acquired using the Leica Application Suite X software using an overlap of 10% per image and a resolution of 1024 x 1024 (0.651 x 0.651 µm per pixel).

Confocal images were acquired with a 63x oil immersion lens (1.4NA, Zeiss), using a Zeiss LSM800 confocal microscope and the ZEN 2.5 Zeiss software package, at a resolution of 2048 x 2048 (0.0495 x 0.0495 µm per pixel).

### Quantification of plaque number and microglia/plaque association

Image analysis was conducted blinded using a semi-automated ImageJ pipeline, where the user draws the outline of the region of the brain in each image to be analysed and inputs Gaussian filter values and thresholds for each channel. For each image, the pipeline then automatically applies a difference-of-Gaussian filter using Clij2 ^47^, followed by automated thresholding and subsequently measures total area and intensity of the selected channels. For individual plaque analysis, the total plaque region of interest (ROI) is split into individual ROIs, then using the ROI Manager, each ROI is given a unique name and subsequently area and intensity are measured for each plaque. For concentric ring analysis, the plaque ROI is enlarged and using logical operations (XOR) the original ROI is subtracted from the enlarged ROI to generate concentric rings with a user defined increase in size around the original selection (here 3 x 10µm). Each concentric ring is given the same name as the original ROI they were generated from + a suffix to denote its increase in size.

For each of these rings and the plaque ROIs, the total ROI size as well as selected channel area and intensity within these ROIs is measured. To quantify the area that a selected channel occupies in the vicinity of each plaque specifically within microglia, a threshold for Iba1 is set to obtain an ROI for the entirety of microglia.

Then, using logical operators with the ROI manager (AND), ROIs corresponding to microglia colocalizing with plaque ROIs and concentric rings are obtained. Lastly, the total ROI size as well as selected channel area and intensity within these ROIs is measured. For each processed image a csv file is created, which were subsequently processed, analysed and plotted using R (4.1.1) and R Studio (2024.09.0+ 375)^48^. For percent area calculations, thresholded signal area was divided by the total ROI area and multiplied by 100.

### 3D evaluation of plaque morphology and microglial clustering around plaques

For the evaluation of plaque volume, sphericity and proximity of microglia and Aβ to plaques, 5 plaques per mouse were picked randomly and 63x confocal z-stacks were acquired along the cortex (z-distance 1.7 µm). Using an automated FIJI script, a 3D difference-of-Gaussian filter was applied and images were made isotropic using Clij2. Then, using the 3D ROI manager ^49^, individual ROIs were imported from each microglia nucleus (based on PU.1^+^ nuclei) or Aβ (3552), from each thiazine^+^ plaque (excluding objects touching the image edges, as well as top and bottom z-slices) and from the total image volume. 3D measurements of each plaque (volume, sphericity, etc), distance of each PU.1^+^ nucleus or Aβ ROI to each plaque and colocalization between each plaque and the total volume of microglia were obtained using 3D manager built-in functions. These measurements were exported as csv files and further processed, analysed and plotted using R (4.1.1) and R Studio (2024.09.0+ 375). Measurements of 5 plaques per mouse were averaged and images where PU.1 ROI separation was not achieved were excluded. Representative 3D isotropic images were made using napari ^50^.

### Protein extraction

Whole hemispheres were lysed following previously published protocol ^51^ and kept at 4°C during all steps. Briefly, hemispheres were lysed in DEA buffer (0.2% diethylamine in 50 mM NaCl, pH 10, and protease inhibitor mix (Sigma, P8340) using the Precellys homogenizer in 2 mL Tissue Homogenizing CKmix tubes (Precellys, P000918-LYSK0-A). Lysate was centrifugated 10 min at 4000g and supernatants were ultracentrifugated at 100 000g before collection. Samples were neutralized by adding 10% of 0.5 M Tris-HCl buffer (pH 6.8) to each sample (DEA fraction). Remaining pellets in Precellys tubes were lysed in RIPA buffer (20 mM Tris-HCl (pH 7.5), 150 mM NaCl, 1 mM Na2EDTA, 1% NP-40, 1% sodium deoxycholate, 0.1% SDS, and protease inhibitor mix. RIPA lysates were centrifuged 10 min at 4000g, and supernatant was ultracentrifugated at 100 000g for 60 min before collection (RIPA fraction). The remaining material in Precellys tubes was resuspended in 70% formic acid with protease inhibitor mix and sonicated for 7min. Samples were centrifuged at 20 000g for 20 min and collected supernatant was diluted 1:20 in pH neutralizing 1M Tris-HCl buffer (pH 9.5) (FA fraction). Protein concentration was measured using Pierce^TM^ Bicinchoninic acid (BCA) assay (Thermo Scientific, 23225).

### Enzyme-linked immunosorbent assay (ELISA)

Aβ levels in FA fraction and CSF were determined using the Meso Scale Discovery (MSD) platform and the V-PLEX Plus Aβ Peptide Panel 1 (6E10) Kit (Meso Scale Discovery, K15200G). FA samples were diluted 1:10 in dilution buffer, CSF was diluted 1:60. Cxcl10/IP-10 levels in DEA fraction were measured using the MSD U-PLEX Mouse IP-10 Assay (Meso Scale Discovery, K152UFK) at a dilution of 1:2.

TREM2 levels in DEA and RIPA fractions, as well as CSF, were measured using the MSD platform as described previously ^52^. Briefly, MSD-gold Streptavidin-coated 96-well plates (Meso Scale Discovery, L15SA-1) were coated in 3% bovine serum albumin (BSA) + 0.05% Tween 20 in PBS (blocking buffer) overnight at 4°C. Sample is diluted in 1% BSA + 0.05% Tween 20 in PBS + protease inhibitor mix (Sigma, P8340), 1:10 for DEA, 1:2 for RIPA and 1:35 for CSF. The plate is incubated with 25 µL/well capture antibody in blocking buffer for 90min, followed by 120 µL/well sample for 2h, detection antibody 50 µL/well for 60 min and SulfoTAG antibody 25 µL/well for 60 min at 600 rpm at RT. In between each incubation the plate is washed 3x with 0.05% Tween 20 in PBS. Before read-out, the plate is washed 2x in PBS, 150 µL/well MSD Read buffer T (Meso Scale Discovery, R92TC-1) is added and is read immediately.

| Capture Antibody | Concentration | Catalogue number | Company |
| --- | --- | --- | --- |
| Goat-anti-Trem2 biotinylated | 0.125 µg/mL (1:800) | BAF1729 | R&D Systems |
| **Detection Antibody** |  |  |  |
| Rat-anti-Trem2 (5F4) | 1 µg/mL (1:1000) | n/a | In house, see <sup>27</sup> |
| Rabbit-anti-Trem2 (HL1738) | n/a (1:10.000) | MA5-31267 | Thermo Fischer |
| **SULFO-TAG Antibody** |  |  |  |
| Goat-anti-rat-SULFO-TAG | 0.5 µg/mL (1:1000) | R32AH-1 | Meso Scale Discovery |
| Goat-anti-rabbit-SULFO-TAG | 0.5 µg/mL (1:1000) | R32AB | Meso Scale Discovery |

### Western blot

4x Laemli Buffer (Biorad, 1610747) + 5% (v/v) β-mercapto-ethanol was added to all samples. For Aβ immunoblotting DEA and FA lysates were loaded on Novex™ WedgeWell™ 10 to 20%, Tris-Tricine, 1.0 mm gels (Thermo Fischer, EC66255) and run in 1x Tris-Tricine-SDS buffer. For Trem2 and APP analysis, samples were run on 12% Tris-Glycine gels in Tris-Glycine-SDS buffer. Protein was transferred to nitrocellulose membrane using wet transfer in Tris-Glycine buffer (pH 7.5).

Membranes were boiled in PBS for 15 min before blocking 1-2 hours in 0.2% I-Block™ Protein-Based Blocking Reagent (Applied Biosystems, T2015) and 0.1% Tween in Tris-buffered saline (TBS) (blocking buffer). Membranes were incubated in primary antibody in blocking buffer O/N at 4°C while shaking. After 3x 10min washes in TBS + 0.05% Tween (TBS-T) membranes were incubated in secondary antibody in blocking buffer for 1 h at RT while shaking. Membranes were developed using Pierce™ ECL Western Blotting-Substrate (Thermo Scientific, 32106) and signals visualized using autoradiographic development using Fuji reed blue film, RX-N (Fuji Film, 47410 19289) or using the Amersham Imagequant 800.

| Primary antibody | Concentration | Catalog number | Company |
| --- | --- | --- | --- |
| Rat-anti-Aβ (2D8) | 1:50 from hybridoma supernatant | n/a | n/a |
| Rat-anti-TREM2 (5F4) | 1 μg/mL (1:1000) | n/a | In house, see <sup>27</sup> |
| Rabbit-anti-APP (Y188) | 0.384 μg/mL (1:1000) | ab32136 | Abcam |

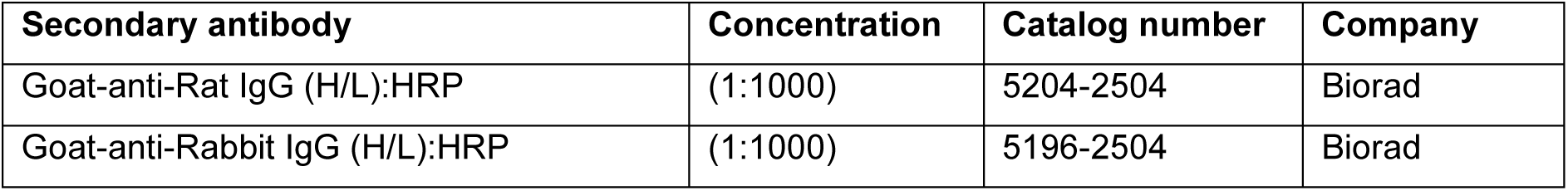

| Secondary antibody | Concentration | Catalog number | Company |
| --- | --- | --- | --- |
| Goat-anti-Rat IgG (H/L):HRP | (1:1000) | 5204-2504 | Biorad |
| Goat-anti-Rabbit IgG (H/L):HRP | (1:1000) | 5196-2504 | Biorad |

### Magnetic-activated microglia sorting (MACS) from mouse brain

Following perfusion, mouse brains were harvested and kept in Hanks’ buffered salt solution with Ca^2+^ and Mg^2+^ (HBSS) (Gibco, 14025092) + 7 mM HEPES (Gibco, 15630080) + 2x GlutaMAX (Gibco, 35050061) on ice. To remove meninges, brains were gently rolled on a clean piece of Whatman paper. Cerebellum, pons and olfactory bulb were removed, two hemispheres were split and any remaining meninges were removed with tweezers using a dissection microscope. Each hemisphere was cut into pieces using a scalpel and brain tissue was dissociated following manufacturer’s instructions using the Neural Tissue Dissociation Kit (P) (Miltenyi, 130-092-628) supplemented with 5 µM Actinomycin D (Cell Signaling Technology, 15021) and 2 µM Anisomycin (Cell Signaling Technology, 2222) in gentleMACS C-tubes (Miltenyi, 130-096-334) using a gentleMACS Dissociator (Miltenyi). Homogenized tissue was run through a 40 μm cell strainer (Corning, 352340) and pelleted by centrifugation. Pellets were resuspended in HBSS with 0.25% fatty acid free BSA (Sigma-Aldrich, A8806), incubated with magnetic Cd11b+ MicroBeads (Miltenyi, 130-093-634) and run twice over MS columns (Miltenyi, 130-042-201). Cells were counted using trypan blue, aliquoted into tubes, centrifuged and snap frozen in liquid nitrogen until further processing.

### Sample preparation for mass spectrometry

CSF samples were prepared as described previously ^45^. Briefly, a volume of 5 µL CSF was used for proteolytic digestion. Proteins were reduced by the addition of 2 µL of 10 mM dithiothreitol (Biozol, Germany) in 50 mM ammonium bicarbonate and incubated for 30 min at 37°C. Cysteine residues were alkylated by the addition of 2 µL 55 mM iodoacetamide (Sigma Aldrich, US) and incubated for 30 min at room temperature in the dark. Afterwards, the reaction was quenched by adding another 2 µL of 10 mM dithiothreitol. Proteolytic digestion was performed using a modified protocol for single-pot solid-phase enhanced sample preparation (SP3) ^53^. After binding proteins to 40 µg of a 1:1 mixture of hydrophilic and hydrophobic magnetic Sera-Mag SpeedBeads (GE Healthcare, US) with a final concentration of 70% acetonitrile for 30 min at room temperature, the beads were washed four times with 200 µL 80% ethanol. For proteolytic digestion, 0.1 µg LysC and 0.1 µg trypsin (Promega, Germany) were added in 20 µL 50 mM ammonium bicarbonate followed by an incubation for 16 h at room temperature. The magnetic beads were retained in a magnetic rack and the supernatants were filtered with 0.22 µm spin filters (Spin-x, Costar) to remove remaining beads and dried by vacuum centrifugation.

### Liquid chromatography tandem mass spectrometry (LC-MS/MS) of CSF

Dried peptides were dissolved in 20 µL 0.1% formic and 5.5 µL were separated on a nanoElute nanoHPLC system (Bruker, Germany) on an in-house packed C18 analytical column (15 cm × 75 µm ID, ReproSil-Pur 120 C18-AQ, 1.9 µm, Dr. Maisch GmbH) using a binary gradient of water and acetonitrile (B) containing 0.1% formic acid at flow rate of 300 nL/min (0 min, 2% B; 2 min, 5% B; 62 min, 24% B; 72 min, 35% B; 75 min, 60% B) and a column temperature of 50°C. The nanoHPLC was online coupled to a TimsTOF pro mass spectrometer (Bruker, Germany) with a CaptiveSpray ion source (Bruker, Germany). A Data Independent Acquisition Parallel Accumulation–Serial Fragmentation (diaPASEF) method was used for spectrum acquisition. Ion accumulation and separation using Trapped Ion Mobility Spectrometry (TIMS) was set to a ramp time of 100 ms. One scan cycle included one TIMS full MS scan with 26 windows with a width of 27 m/z covering a m/z range of 350-1001 m/z. Two windows were recorded per PASEF scan. This resulted in a cycle time of 1.4 s.

### Mass spectrometry data analysis

The software DIA-NN version 1.8.1 was used to analyse the data ^54^. The raw data was searched against a one protein per gene database from Mus musculus (UniProt, 21709 entries, download: 2024-02-19) combined with a database of common human contaminations (123 entries) using a library free search. Trypsin was defined as protease and two missed cleavages were allowed. Oxidation of methionines and acetylation of protein N-termini were defined as variable modifications, whereas carbamidomethylation of cysteines was defined as fixed modification. The precursor and fragment ion m/z ranges were limited from 350 to 1001 and 200 to 1700, respectively. An FDR threshold of 1% was applied for peptide and protein identifications. The mass accuracy and ion mobility windows were automatically adjusted by the software. The match between runs option was enabled.

The statistical analysis was performed with the software Perseus version 1.6.2.3 ^55^. First, a one-way ANOVA was used to determine statistically significant differences between the means of the groups. Afterwards, individual Student’s *t*-tests were applied to evaluate proteins with a significant abundance difference between 1, 3, and 10 mg/kg anti-Aβ compared to isotype control treatment. Additionally, isotype control samples were compared with sample from three months old untreated mice. A permutation based false discovery rate estimation was used with a FDR of 5% at s0 = 0.1 as threshold ^56^.

### RNA isolation, RT–qPCR, and library preparation

To prepare for RNA-seq, approximately 100,000 CD11b+ microglia isolated by MACS were used for RNA extraction by the RNeasy Plus Micro Kit (Qiagen, #74034). The extracted RNA was then resuspended in nuclease-free water for RNA-seq library preparation. Libraries for 30 total RNA samples were prepared using the Lexogen QuantSeq 3′ mRNA-Seq V2 Library Prep Kit FWD with Unique Dual Indices (Lexogen 193.384) and the UMI Second Strand Synthesis Module, following the manufacturer’s protocol to identify and remove PCR duplicates. In brief, total RNA was used as input for oligo(dT) priming during reverse transcription, followed by RNA removal. Unique Molecular Identifiers (UMIs) were incorporated during second-strand synthesis. The cDNA was purified using magnetic beads, amplified with 18 cycles of PCR, and subsequently purified again. Library quantity and quality were assessed using a TapeStation D1000 ScreenTape (Agilent 5067-5582). Equimolar pooling of libraries was performed, and sequencing reads were generated on one lane of an Illumina NovaSeq X 10B cartridge (75 bp single-end) by SeqMatic (Fremont, CA, USA).

### RNA-seq data analysis

RNA-seq data was processed using nf-core/rnaseq v3.11.2 (https://doi.org/10.5281/zenodo.1400710) of the nf-core collection of workflows ^57^. Reads were aligned to the GRCm39 release of the mouse genome, and gene annotations were obtained from Gencode M31. To account for the use of unique molecular identifiers (UMIs) in the library preparation protocol, the following arguments were passed to the STAR aligner (version 2.7.9a, ^58^): --alignIntronMax 1000000 --alignIntronMin 20 --alignMatesGapMax 1000000 --alignSJoverhangMin 8 --outFilterMismatchNmax 999 --outFilterType BySJout -- outFilterMismatchNoverLmax 0.1 --clip3pAdapterSeq AAAAAAAA. After alignment, UMIs were extracted with the following regular expression: ^(? *P*. {6})(? *P*. {4}).∗ . As each transcript is only represented by a single sequence, the --noLengthCorrection parameter was passed to the salmon (version 1.10.1, ^59^) gene-level quantitation step. The pipeline was executed with Nextflow v23.10.0 ^60^. Downstream analysis was performed using R (version 4.4.0) using the limma/voom workflow ^61^ to fit linear models for each quantifiable gene. Library sizes were estimated using the TMM method ^62^, and we fit a linear model with treatment group and sex as fixed covariates, and takedown-batch as a random effect with the voomLmFit function from the edgeR R package (version 4.2.0, ^61^). Sample weights were included by setting the sample.weights argument to TRUE. Differentially expressed genes were identified with the eBayes function from the limma R package (version 3.60.0, ^63^), setting the robust=TRUE argument. P-values were corrected for multiple-testing according to ^64^. Gene set enrichment analyses were performed with the fgsea R package (version 1.30.0, ^65^), with gene sets obtained via the msigdbr package (7.5.1, doi: 10.32614/CRAN.package.msigdbr).

### Lipid Extraction

Cell pellets (100,000 MACS-sorted cells) were suspended in 400 μL of a 3:1 (v/v) butanol/methanol extraction buffer with stable isotope-labeled internal standards and mixed for 5 min at 600 rpm on a plate shaker at room temperature. Plates were stored for one hour at -20°C and centrifuged at 21 000g for 5 minutes at 4°C. After centrifugation, 200 μL of the supernatant was collected and dried under a continuous stream of nitrogen gas. The dried extracts were reconstituted in 200 μL of LCMS-grade methanol for subsequent analysis.

### LC-MS Analysis of Lipids

Lipid analysis was performed using an Agilent Infinity II 1290 UHPLC coupled with a QTRAP 6500+ mass spectrometer. Lipids were analysed in both positive and negative ionization modes and resolved on a UPLC BEH C18 column (150 × 2.1 mm, 1.7 μm, Waters Corp.) at 55°C with a 0.25 mL/min flow rate, following the buffer and gradient schedule from Logan et al. (2021). Data acquisition, peak integration, and quantification were conducted using MultiQuant (version 3.3, ABSciex) with a minimum signal-to-noise ratio of 5 and at least 8 points across the baseline.

### Statistical analysis

Unless indicated otherwise in the methods, statistical analysis was performed in R studio (R version 4.2.3) ^48^. Data are shown with the mean standard error of the mean (±SEM) unless indicated otherwise. For normally distributed data a one-way ANOVA was applied. Statistical evaluations are displayed as follows: *P < 0.05; **P < 0.01; ***P < 0.001; ****P < 0.0001. Graphs were plotted using the *tidyverse* package and statistical significance was plotted using the *ggsignif* package ^66,67^.

## Results

### Chronic anti-Aβ treatment reduces amyloid with greater preference for loosely aggregated fibrils in a dose-dependent manner

APP-SAA KI mice have previously been reported to develop MX-04 positive plaques with an increased brain Aβ_42/40_ ratio compared to wild-type mice at 4 months of age^35^. To investigate the progression of pathology in APP-SAA KI mice and determine the optimal treatment window for early intervention, we investigated amyloid plaque deposition at 3, 6 and 12 months of age. Immunostaining of brain sections with thiazine red, a dye that binds to core plaques ^68^, and an Aβ-antibody (3552), which binds to total Aβ ^69^, indicates that APP-SAA KI mice already show the sporadic appearance of loosely aggregated Aβ fibrils in plaque-like structures as early as 3 months (Fig. S1A,B). Furthermore, these diffuse plaques are already surrounded by Lamp1-positive neuritic dystrophy. Thiazine-positive dense core plaques appear at 6 and 12 months together with Trem2 positive microglia (Fig. S1C, D). Moreover, Trem2 protein levels increase with age and amyloid plaque accumulation, suggesting a strong induction of DAM activation concomitant with the accumulation of plaques, as observed in other amyloidogenic mouse models ^70,71^ (Fig. S1E). Interestingly, microglia also appear to cluster around diffuse plaques already starting at 3 months (Fig. S1F). Due to the Austrian mutation ^72^, plaques of APP-SAA KI mice contain a large amount of Aβ38, which is not typically incorporated into amyloid plaques in sporadic AD patients ^73,74^. Most Aβ accumulates in the insoluble (FA) fraction, with limited detection in the soluble (DEA) fraction (Fig. S1G, H). Importantly, the anti-Aβ antibody used for dosing studies in this model (Aducanumab) binds to residues 3-7 of Aβ, which does not overlap with the three dominant mutations inserted into the APP gene (Fig. S1I) ^75^.

To understand the dose-dependent effects of a chronic intervention treatment paradigm and potential dose-dependent effects, APP-SAA KI mice were treated with 1, 3 or 10 mg/kg of anti-Aβ or an isotype antibody (1 or 10 mg/kg) weekly i.p. for 16 weeks (Fig. 1A). To simulate an early intervention paradigm where plaques already start to form, a time-point of 4.5 months was chosen to start treatment. The [18]F-florbetaben (FBB)-PET ^76^ signal detected in 8.5 months old APP-SAA KI mice is reduced in a dose-dependent manner by anti-Aβ treatment (Fig. 1B, C). The reduction of Aβ was confirmed by Western Blot analysis of the formic acid (FA) fraction (Fig. 1D). In contrast, sAPP (Fig. 1D), full length APP and its C-terminal fragments (CTF) were not altered in total brain extracts by the antibody treatment, confirming no changes in APP processing upon treatment (Fig. S2A).

**Figure 1:**
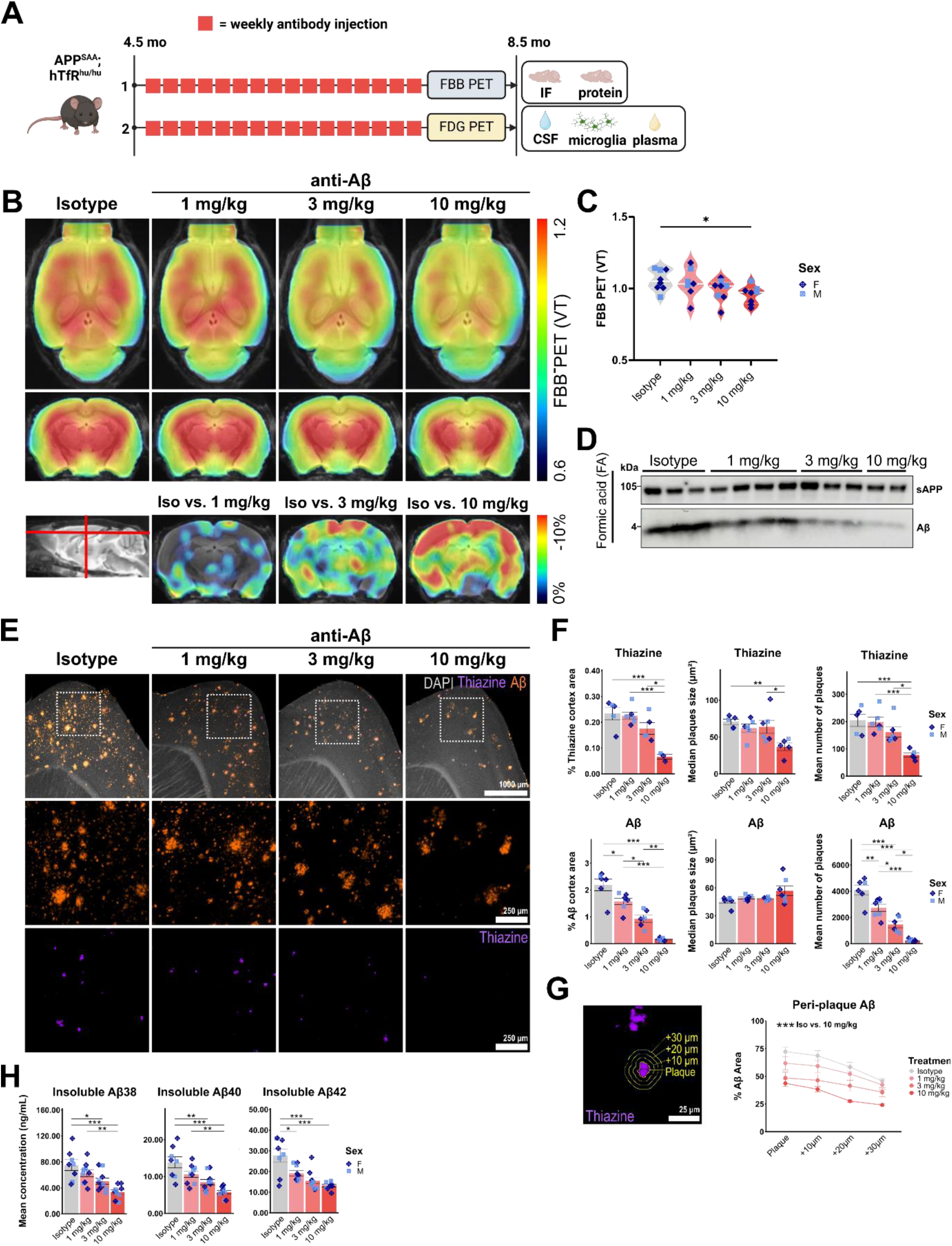
Chronic anti-Aβ treatment reduces Aβ levels in a dose-dependent manner. **(A)** Schematic of the study design and collected materials of anti-Aβ or isotype (4D5) antibody treatment cohorts. (**B**) Axial and coronal FBB-PET standardized uptake value (VT), and coronal FBB-PET (% change from Isotype) per group projected upon a standard magnetic resonance imaging (MRI) T1 atlas. (**C**) Quantification of FBB-PET (VT). (**D**) Western blot showing soluble (s)APP and Aβ in the formic acid extracted fraction. (**E**) Representative immunofluorescent images of sagittal cortical sections showing DAPI (grey), thiazine (purple) and Aβ (3552 antibody, orange) with insets showing thiazine and Aβ. (**F**) Quantification of % cortical plaque and Aβ area, size and number. (**G**) Example of concentric plaque regions of interest (ROIs) and quantification of Aβ (3552) signal in these ROIs. (**H**) ELISA quantification of formic acid extracted insoluble Aβ. *: P < 0.05; **: P < 0.01; ***: P < 0.001. One-way ANOVA with Tukey’s post hoc test (C, F, G, H). Schematic (A) was created with BioRender.com.

Immunostaining of brain sections with thiazine red and Aβ-antibody (3552), indicates that plaques and fibrils are mainly located in the cortex (Fig. 1E), hippocampus and cortico-amygdala area (COA) (Fig. S2B). Treatment with anti-Aβ lowers both core plaques stained by thiazine and total Aβ levels in a dose dependent manner in all three brain regions (Fig. 1F and S2C). Interestingly, percent total Aβ area and number was already reduced with 1 mg/kg anti-Aβ treatment, as shown by immunostaining with a pan-Aβ antibody, as opposed to a reduction in thiazine-positive, dense-core plaque area and number only upon treatment with 10 mg/kg anti-Aβ. This suggests that loosely packed Aβ fibrils are more readily removed and/or prevented from aggregating into plaques. Analysis of percent area covered by Aβ at and around dense core plaques in concentric rings indicated a dose-dependent reduction in peri-plaque Aβ (Fig. 1G). To investigate whether this could be associated with changes in plaque compactness we performed confocal imaging of whole plaques and analysis of 3D reconstructed images. However, no changes were observed in plaque compactness when quantifying the Aβ/Thiazine volume ratio or mean distance of Aβ objects to thiazine (Fig. S2D, E). Lastly, whole brain levels of different Aβ species were quantified using an Aβ-triplex ELISA, which shows a significant dose-dependent reduction at 3 and 10 mg/kg doses in Aβ_38_, Aβ_40_ and Aβ_42_ levels (Fig. 1H). In parallel in CSF, although individual Aβ_38_, Aβ_40_ and Aβ_42_ levels showed high between-mouse variability, an increase in the CSF Aβ_42/40_ ratio was observed (Fig. S2F). Overall, these results indicate a significant removal of Aβ and prevention of plaque formation in APP-SAA mice upon chronic anti-Aβ treatment, with a higher efficiency for loosely aggregated forms of Aβ, in a dose-dependent manner.

### Chronic anti-Aβ treatment does not worsen microhaemorrhages in APP-SAA KI mice

Clinical use of anti-Aβ antibodies is associated with an increased risk of microbleeds, the cause of which has been hypothesized to be linked to the treatment induced-immune response, as well as amyloid deposition in cerebral blood vessels^11,12^. To investigate vascular amyloid pathology in the APP-SAA KI mouse, we used HS169, a compound that binds to both diffuse and fibrillar plaques and vascular deposits ^77^. We found that APP-SAA KI mice show cerebral amyloid-angiopathy (CAA)-like Aβ aggregation in meningeal and cerebral blood vessels at the age of 8.5 months (Fig. S3A). By quantifying HS169 area, we confirmed a dose-dependent reduction in total cortical Aβ, but observed no significant changes in HS169 co-localizing with vascular marker laminin (α1) upon treatment (Fig. S3B, C). Previous reports also described that anti-Aβ antibody treatment in mouse models can lead to an increase in microbleeds ^30,33^. To investigate whether chronic anti-Aβ treatment in APP-SAA KI mice is associated with microbleeds in a dose-dependent manner, Prussian blue staining of hemosiderin was analysed. Surprisingly, evidence of past microbleeds, in the forms of Prussian blue-positive microglial cell shaped foci, was observed in sections of all treatment conditions including treatment with the isotype control (Fig. S3D). However, no significant changes were observed in foci count or percent area across sections with anti-Aβ treatment (Fig. S3E). To confirm whether this effect could be associated with antibody treatment in general, control sections of untreated 3, 6 and 12-month-old APP-SAA KI mice were analysed (Fig. S3F). Interestingly, these mice also show Prussian blue foci without antibody treatment at 6 and 12 months, but not at 3 months (Fig. S3F). These results suggest that the APP-SAA KI mouse model develops spontaneous microbleeds, which may be related to the presence of CAA, both of which appear not worsened by antibody treatment as measured by Prussian blue staining.

### CSF proteome changes indicate reduced immune activation and neuropathology after chronic anti-Aβ treatment

CSF biomarkers are routinely used in clinical practice to diagnose AD ^78^. In addition, antibody treatments in clinical trials help to identify novel biomarkers that can predict better or worse disease outcomes and could potentially be used to monitor target engagement upon anti-amyloid antibody treatment in patients ^79^. To investigate potential CSF biomarkers of anti-Aβ treatment, we analysed terminally collected CSF of treated mice using liquid chromatography tandem mass-spectrometry (LC-MS/MS) using shotgun proteomics with label-free quantification. CSF from 3-month-old APP-SAA mice was included as a baseline/pre-treatment comparison. When comparing 8.5-month-old isotype treated mice to 3-month-old APP-SAA KI mice, several inflammatory marker proteins such as Trem2, Lag3, Lyz2, and Sod2 as well as the neurodegenerative marker Tau (Mapt) showed an increased abundance, which was partially attenuated with 10 mg/kg anti-Aβ treatment (Fig S4A, B).

Comparing individual treatment doses to isotype control, several disease-related proteins showed decreased abundances, which however were not significant after FDR correction (Fig. 2A, B). The immune cell-related proteins Cst7, Cd84, Ctsz and Trem2 showed a dose-dependent response (Fig. 2C). ELISA of CSF showed a sex-specific effect with Trem2 levels decreasing significantly only in males (Fig S4C, D). In terminally collected plasma no significant treatment dependent differences in sTREM2 concentrations were observed (Fig S4E). Interestingly, LC-MS/MS analysis also identified neuronal proteins in CSF, such as Tau, Camk2a, α-Synuclein (Snca) and Gap43, which showed a dose-dependent rescue, suggesting reduced neuritic pathology upon anti-Aβ treatment (Fig. 2D). To confirm if the reduction of neuronal markers reflects reduced neuropathology, neuritic dystrophy was analysed using Lamp1 immunofluorescence. In concurrence with the reduced number of thiazine positive plaques and the reduction of neuronal markers, a reduction of Lamp1 neurite dystrophy was observed at 10 mg/kg anti-Aβ (Fig 2E, F). In addition, we found at 10 mg/kg a significant reduction in the peri-plaque Lamp1 signal (Fig 2G), suggesting that residual plaques induce less neurite dystrophy in their immediate surrounding. In summary, CSF proteomics detects both neurite pathology and immune cell-related markers, which respond in a dose-dependent manner to anti-Aβ treatment.

**Figure 2:**
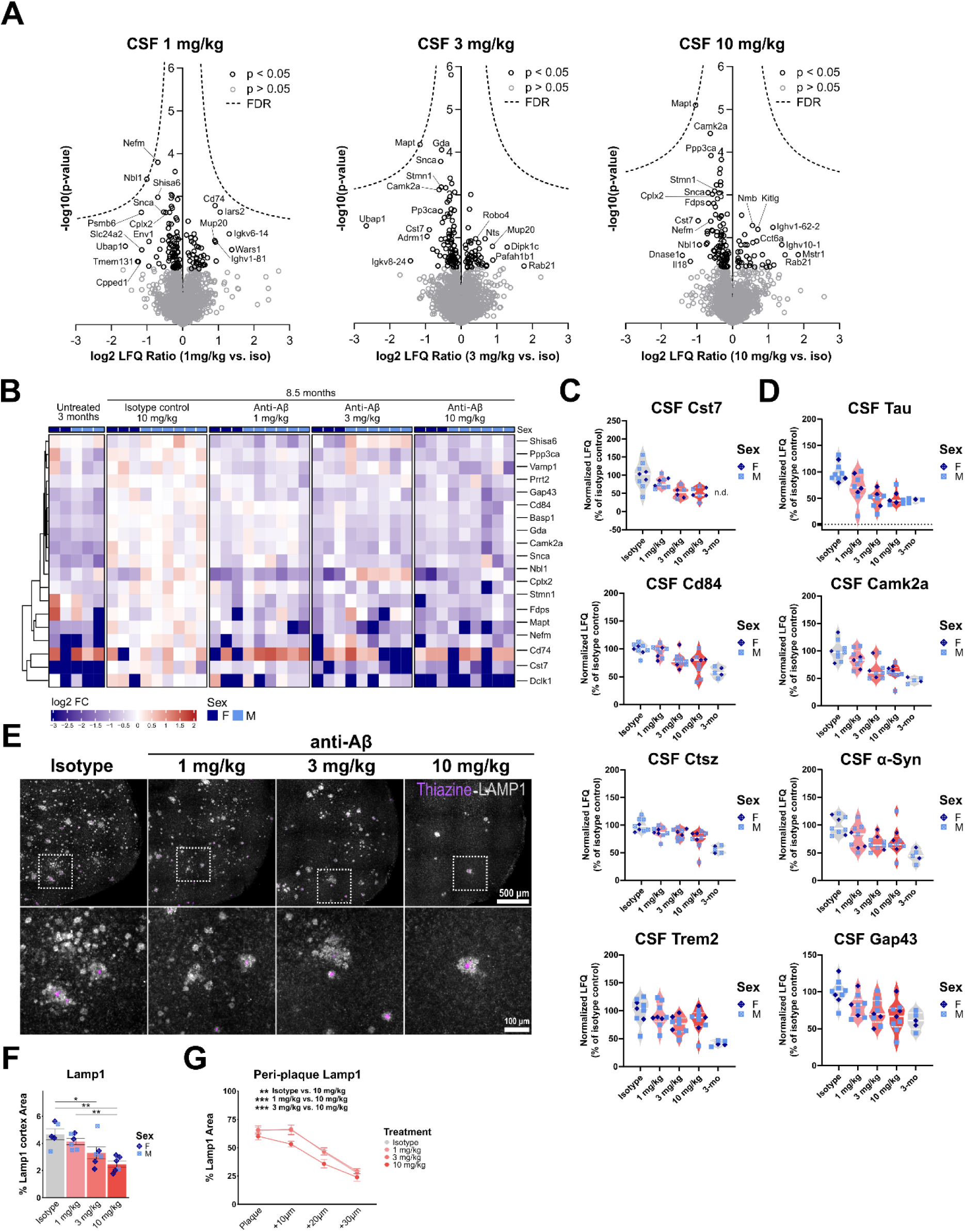
CSF proteome changes indicate reduced neuropathology and immune activation after chronic anti-Aβ treatment. **(A)** Volcano plots showing upregulated and downregulated proteins in CSF. Each volcano shows changes in comparison to isotype treated mice. (**B**) Heatmap showing unbiased clustering of top 20 changed proteins relative to isotype control treated mice (cut-off p<0.05, log2FC < -0.5 or >0.5). (**C**) Normalized LFQ plots of microglial and (**D**) neuronal proteins relative to isotype. (**E**) Representative fluorescent images showing thiazine (purple) and Lamp1 (grey) in sagittal cortical sections. (**F**) Quantification of Lamp1 % area in the cortex. (**G**) Quantification ofLamp1 in concentric rings in and around thiazine plaques. **: P < 0.01; ***: P < 0.001; ****: P < 0.0001. One-way ANOVA with Tukey’s post hoc test (F, G).

### Chronic anti-Aβ treatment partially returns microglia to a pre-disease transcriptional and lipidomic signature

To investigate how the observed immune cell-related protein changes in CSF are associated with microglial phenotypes in the brain, bulk 3’-RNA sequencing was performed on Cd11b^+^ MACS-sorted microglia. Microglia sorted from 4-month-old APP-SAA KI mice were included as a baseline/pre-treatment comparison. With aging, in all treatment groups and isotype control, a strong increase in microglial activation is observed, with ∼4500 differentially expressed genes (DEGs, false discovery rate [FDR] < 5%) (Fig S5A). These changes correlate to changes identified by Xia *et al.* 2022, between 8-month-old APP-SAA KI mice compared to WT, validating the 4-months’ time-point as a pre-disease control (Fig S5B). Gene set enrichment analysis (GSEA) confirms an enrichment in gene sets associated with metabolic pathways, including oxidative phosphorylation, glycolysis, MTORC1 signaling and cholesterol homeostasis, which show predominantly increased gene expression with age (Fig S5C, D). The gene expression signatures observed are in line with previously defined microglial states in AD ^80^, showing a strong increase in DAM, interferon (IFN), major histocompatibility complex (MHC) and proliferation associated gene expression and a reduction in homeostatic gene expression (Fig S5E), confirming that between 4 and 8 months of age a strong induction of AD associated microglial signatures is induced in APP-SAA KI mice.

When comparing anti-Aβ treatment to isotype, polynomial modelling identified a linear association between anti-Aβ dose and gene expression, detecting ∼400 DEGs (FDR < 5%). Predominantly, a dose dependent attenuation of genes induced during disease progression is observed with anti-Aβ treatment compared to 4-month-old mice at baseline (Fig. 3A). Gene set enrichment analysis of these changes reveals an enrichment in genes associated with cholesterol homeostasis and glycolysis, which are predominantly driven by genes that are reduced in expression upon treatment, as well as complement and interferon signaling, which are predominantly driven by genes that are increased in expression compared to isotype (Fig. 3B, C).

**Figure 3:**
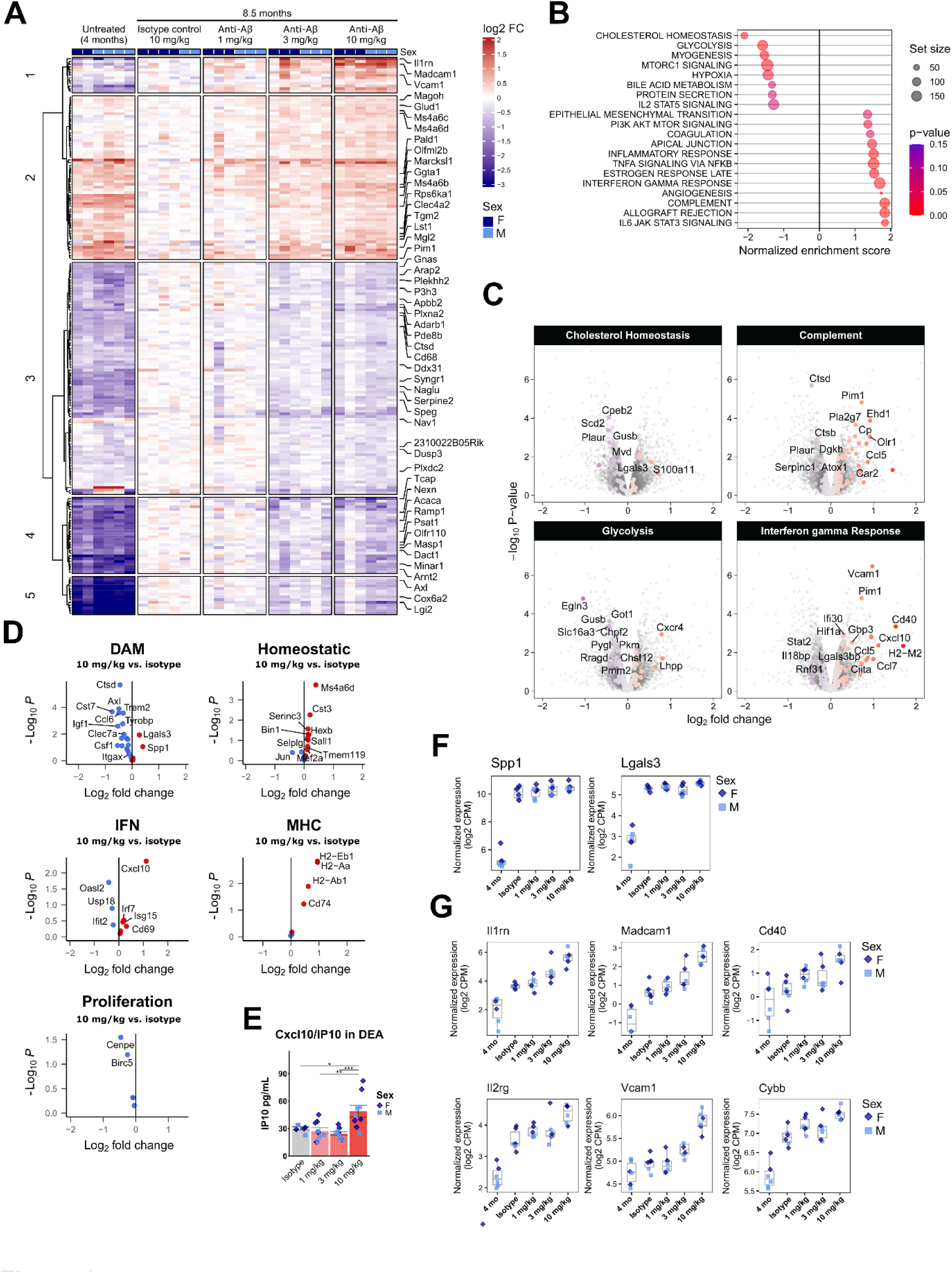
Chronic anti-Aβ treatment partially returns microglia to a pre-disease transcriptional signature. **(A)** Heatmap showing the top 200 differentially expressed genes relative to isotype treated mice. (**B**) Gene set enrichment analysis (GSEA). (**C**) Volcano plots showing differentially expressed genes related to GSEA pathways. (**D**) Volcano plots showing differentially expressed genes related to microglial states from Chen et al. 2021. (**E**) ELISA quantification of Cxcl10/IP10 protein in DEA fraction. (**F**) Boxplots showing gene expression changes in DAM genes *Spp1* and *Lgals3*. (**G**) Boxplots showing examples of genes that increase expression upon anti-Aβ treatment. CPM = counts per million. *: P < 0.05; **: P < 0.01; ***: P < 0.001. One-way ANOVA with Tukey’s post hoc test (E).

When taking a closer look at gene sets associated with known microglial states, we observe a decrease in DAM associated genes such as *Axl*, *Ctsd*, *Cst7*, *Trem2*, *Tyrobp* and *Clec7a*, and an increase in homeostatic genes, such as *Ms4a6d*, *Cst3*, *Sall1*, *Hexb* and *Tmem119* (Fig. 3D). In addition, certain MHC- and IFN-associated genes show a further increase upon anti-Aβ treatment, such as Cxcl10, of which the encoding protein IP10 was found to be increased in DEA soluble brain extract at 10 mg/kg (Fig. 3D). Interestingly, DAM associated genes *Lgals3* (encoding galectin-3) and *Spp1* (encoding osteopontin) did not show a dose-response to treatment, suggesting that anti-Aβ treatment does not completely attenuate the DAM signature (Fig. 3F). The six genes that show the strongest induction upon anti-Aβ treatment, include *Cd40* (antigen presentation)*, Cybb* (Nox2, oxidative burst), *Vcam1* and *Madcam1* (cell adhesion), *Il2rg* and *Il1rn* (cytokine signaling) (Fig. 4G). Overall, these results suggest that anti-Aβ treatment shifts microglia back to a homeostatic state, but also induces a treatment-related inflammatory response, associated with increased antigen presentation, cell adhesion and cytokine signaling, as well as DAM genes *Lgals3* and *Spp1*.

**Figure 4:**
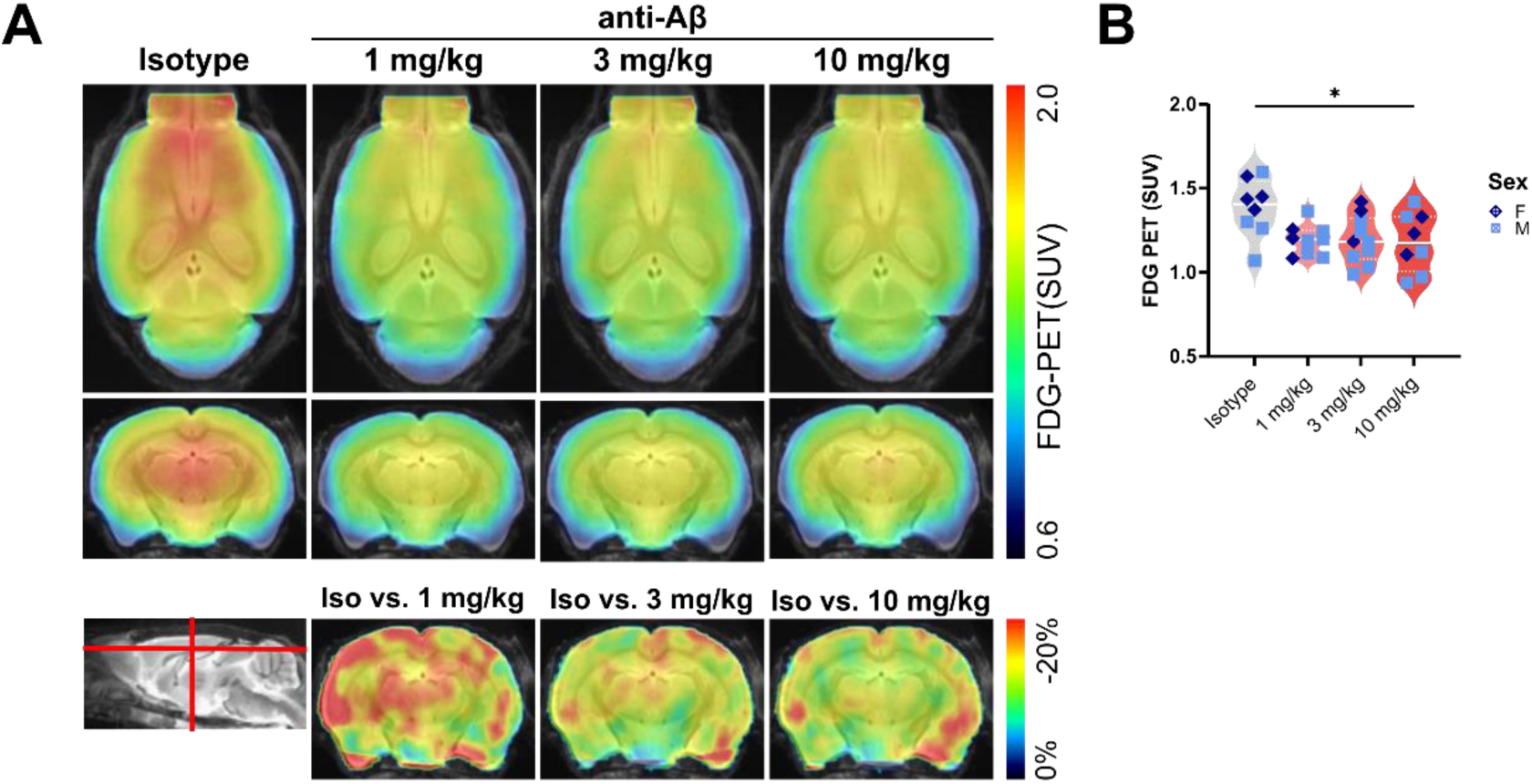
Chronic anti-Aβ treatment decreases microglial glucose uptake. **(A)** Axial and coronal FDG-PET (standardized uptake value (SUV)), and coronal FDG-PET (% change from isotype) per group projected upon a standard magnetic resonance imaging (MRI) T1 atlas. (**B**) Quantification of FDG-PET. *: P < 0.05, One-way ANOVA with Tukey’s post hoc test (B).

To investigate whether alterations in gene expression are associated with changes in microglial lipidome, we performed lipidomic analysis of sorted microglia. Changes in lipid abundance were observed in microglia with aging (Fig. S6A, B).

Although no FDR corrected significant changes were detected upon anti-Aβ treatment, an overall attenuation with treatment towards untreated baseline was observed (Fig.S6C). Most notably, ganglioside GM3 showed a dose-dependent response to anti-Aβ, whereas cholesterol ester (CE) remained high regardless of treatment (Fig.S6D). Interestingly, GM3 was previously shown to be associated with MX-04^+^ microglia in the APP-SAA KI mouse model, whereas CE was observed to be lower in MX-04^+ 35^ (Fig.S6E). These data suggest that GM3 is increased with aging and Aβ pathology in the APP-SAA KI mouse model, whereas CE is increased with aging, but less so associated with Aβ pathology. Taken together, these results indicate that anti-Aβ treatment does not worsen lipid burden in microglia, but mildly attenuates Aβ-induced lipid changes towards pre-treatment baseline and most predominantly the Aβ pathology-associated lipid GM3.

### Chronic anti-Aβ treatment decreases global microglial FDG-uptake and Trem2, but DAM activation around residual plaques remains increased

Bulk-RNA-seq identified a reduction in overall microglial DAM activation and glycolysis with anti-Aβ treatment. Microglial activation in response to Aβ has previously been shown to correlate with a significant increase glucose uptake, which is driven by Trem2 expression and further increased upon Trem2 agonism ^37,42,81,82^. Based on cell-sorting experiments in mouse models of amyloidogenesis, [¹⁸F]-fluorodeoxyglucose (FDG) PET can be used as a readout of microglial glycolytic activity ^42^. To confirm whether the reduced expression of glycolysis-associated genes upon anti-Aβ treatment translates to reduced microglial glucose uptake, we measured FDG-PET after 16 weeks of antibody treatment. Similar to FBB-PET, a dose-dependent reduction in FDG uptake was observed (Fig 4A, B). In parallel, we found a dose-dependent reduction in brain Trem2 protein levels in both DEA and RIPA soluble fractions (Fig. 5A). Trem2 levels correlated with Aβ_42_ levels (Fig. 5B), confirming that the Trem2-induced microglial response is closely linked to the degree of Aβ accumulation. However, as plaques are not completely eliminated with anti-Aβ treatment, we were interested to see whether microglial DAM activation is still induced at residual plaques. Immunostaining confirmed reduced total cortical Trem2 and Cd68 area at 10 mg/kg (Fig. 5C, D), but using concentric plaque analysis we found that treatment did not affect induction of these proteins around residual plaques. ApoE followed a similar pattern, with reduced total cortical area at 10 mg/kg, but was not significantly changed in the plaque penumbra (Fig. S7A, B). In contrast, homeostatic marker P2ry12 was not reduced in total cortex coverage, but also not around plaques (Fig. S7C, D). Taken together, these findings suggest that microglia at residual plaques do not lose their Aβ-induced DAM activation and appear to still be actively dealing with plaques after 16 weeks of anti-Aβ treatment.

**Figure 5:**
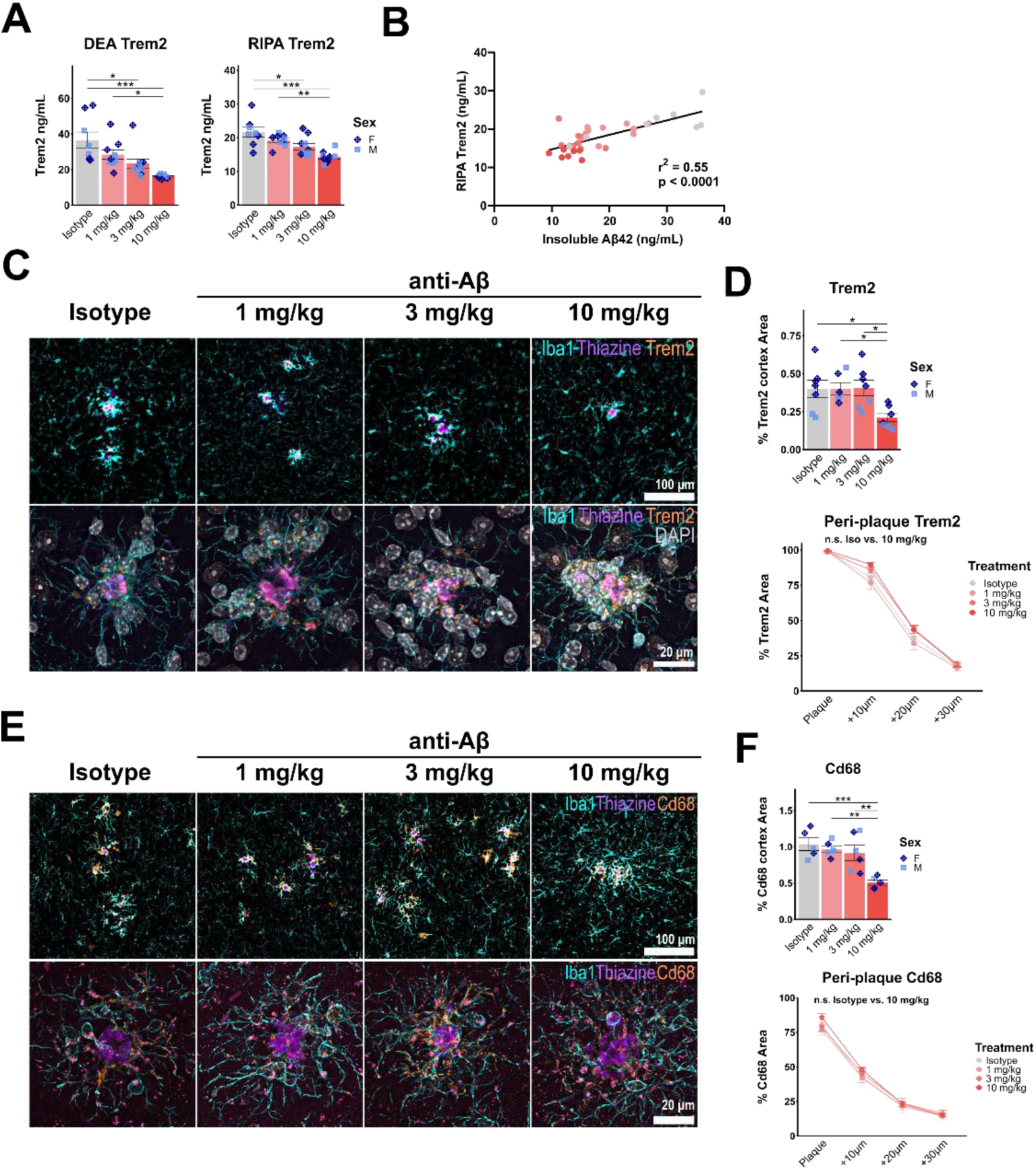
DAM activation is reduced brain-wide, but not around residual plaques. **(A)** ELISA quantification of Trem2 protein in DEA and RIPA brain lysate. (**B**) Correlation between RIPA Trem2 and FA Aβ_42_. (**C**) Representative epifluorescence and confocal images showing dapi (grey), Iba1 (cyan), thiazine (purple) and Trem2 (orange). (**D**) Quantification of percent cortical and concentric plaque-analysis of Trem2. (**E**) Representative epifluorescence and confocal images showing Iba1 (cyan), thiazine (purple) and Cd68 (orange). (**F**) Quantification of percent cortical and concentric plaque-analysis of Cd68. *: P < 0.05; **: P < 0.01; ***: P < 0.001; ****: P < 0.0001. One-way ANOVA with Tukey’s post hoc test (A, D, F), Pearson r correlation (B).

### Chronic anti-Aβ treatment increases microglial clustering and an antibody-induced microglial phenotype around residual plaques

Previous studies have consistently reported increased microglial clustering around plaques upon anti-Aβ treatment in mice and patients ^29,31,83^. However, using concentric plaque analysis we found no changes in peri-plaque Iba1 (Fig. S8A). It is possible that due to the thickness of the brain sections (50 µm), we were unable to distinguish individual microglia based on Iba1 signal with epifluorescence. Therefore, we employed a more sensitive analysis by performing confocal imaging of entire plaques and co-stained with PU.1, a microglial-specific transcription factor. Using 3D reconstruction of PU.1^+^ nuclei and plaques, we could confirm a dose-dependent increase in microglia localized with cell bodies in close proximity (within 10 µm) to plaques (Fig. 6A, B). In addition to microglia, it was recently suggested that acute anti-Aβ treatment can increase astrogliosis, with a trend towards increased Gfap around plaques ^29^. In contrast to these findings, we found no changes in total cortex area coverage of astrogliosis marker Gfap after chronic anti-Aβ treatment, however we did observe increased Gfap coverage at 10 mg/kg around residual plaques (Fig. S8B, C), suggesting that treatment can potentially induce both peri-plaque microgliosis and astrogliosis.

**Figure 6:**
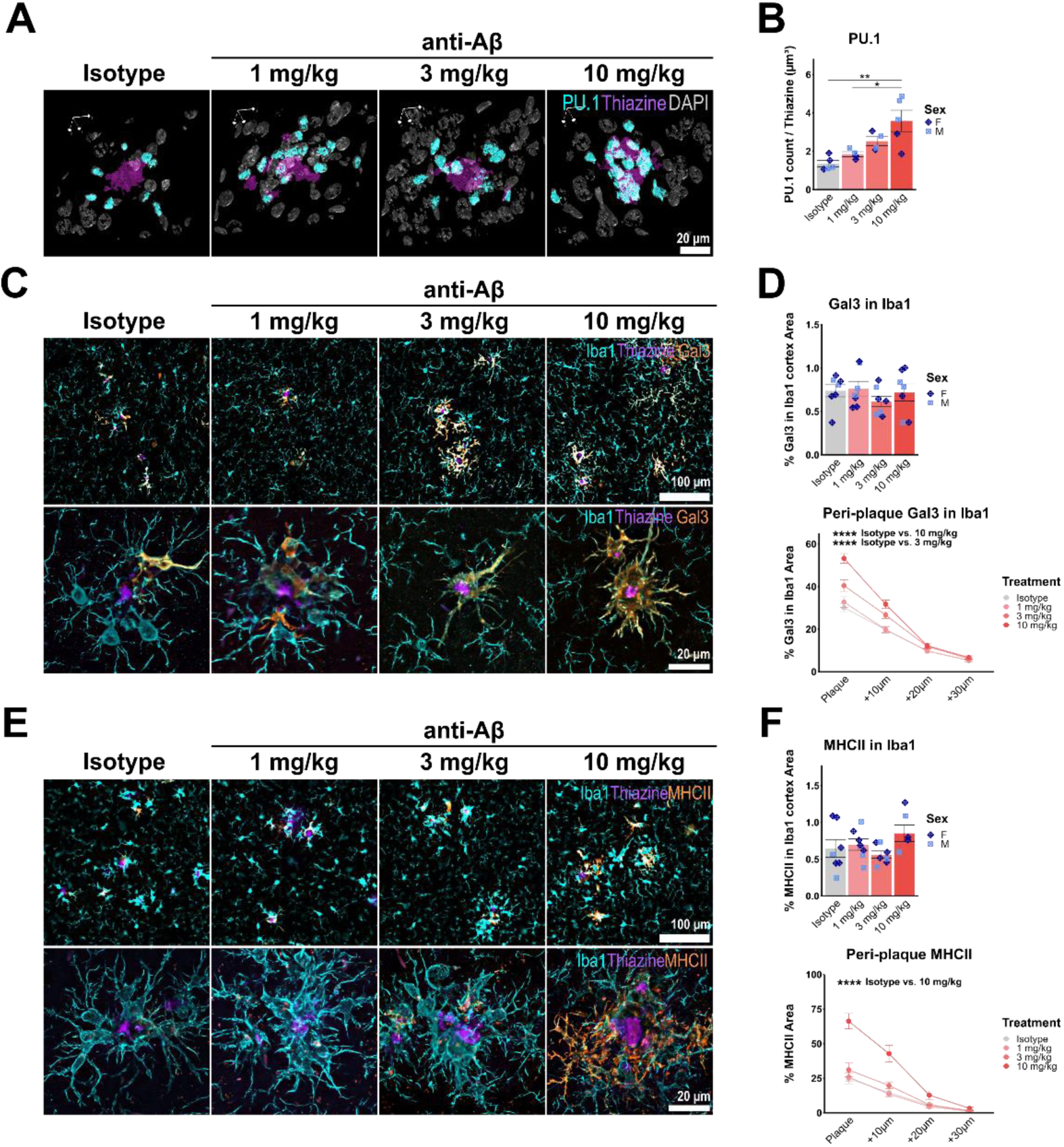
Increased microglial clustering is associated with increased anti-Aβ-induced MHC-II and Galectin-3 around residual plaques. (**A, B**) Isotropic 3D rendering of confocal immunofluorescent images showing thiazine (purple), DAPI (grey) and PU.1 (cyan) and quantification of Spi1^+^ DAPI nuclei number to normalized by plaque volume (**C**) Representative epifluorescence and confocal images showing Iba1 (cyan), thiazine (purple) and co-staining with Galectin-3 (orange). (**D**) Quantification of Galectin-3 in Iba1 in total cortex and in concentric rings around thiazine plaques (**E**) Representative epifluorescence and confocal images showing Iba1 (cyan), thiazine (purple) and co-staining with MHC-II (orange). (**F**) Quantification of MHC-II in Iba1 in total cortex and in concentric rings around thiazine plaques. *: P < 0.05; ***: P < 0.001; ****: P < 0.0001, One-way ANOVA with Tukey’s post hoc test (B, D, F).

To further investigate antibody-treatment induced effects specifically around residual plaques, we stained for Galectin-3 (Fig. 6C, D) and MHC-II (Fig. 5E, F), which we showed to be increased in expression in microglia after anti-Aβ treatment. Despite the finding that overall Galectin-3 and MHC-II coverage in microglia were not different between treatment groups in the cortex using immunofluorescence, a clear dose-dependent increase in protein levels was observed in concentric rings around amyloid plaques for both proteins (Fig. 6D, F). Taken together, these findings indicate increased microglial clustering around plaques, with the concomitant induction of a peri-plaque microglial state that is associated with antigen presentation, phagocytosis and microglial recruitment to plaques.

## Discussion

The current study demonstrates that anti-Aβ treatment results in a dose-dependent removal of Aβ and prevention of plaque formation in APP-SAA KI mice upon chronic treatment. Concomitantly, microglial DAM activation is decreased in a manner that correlates with the reduction in plaque load. However, microglial clustering around residual plaques is increased and these cells display a unique antibody-driven microglial phenotype.

The dose-dependent reduction of amyloid observed in APP-SAA KI mice is in line with the seminal work from Sevigny et al. 2016, who also reported a dose dependent removal of amyloid plaques in mice and humans ^31^. Aducanumab is an antibody that has a strong preference for Aβ fibrils over protofibrils and monomers and should therefore be most efficient at removing aggregated Aβ ^84,85^. Plaques in the APP-SAA KI mouse contain relatively small dense cores and a large amount of loosely aggregated fibrillar Aβ, which was more efficiently removed by anti-Aβ treatment. A significant reduction in FBB-PET was only observed when the dense core plaque number was significantly reduced at the highest dose. This is in line with previous findings that although diffuse plaques bind FBB, dense cores plaques contribute to most of the FBB PET signal ^69^. Interestingly, despite starting at an early disease time-point where only very few plaques should be present, chronic treatment with anti-Aβ was not able to remove nor prevent build-up of plaques completely. This is in line with a recent study that finds that chronic Aducanumab treatment does not remove plaques, but keeps plaque load at a level comparable to the start of treatment ^33^. It is possible that plaques that are already seeded at the start of treatment are not targeted and removed by anti-Aβ as efficiently as loosely aggregated or oligomeric forms of Aβ. How these findings relate to the removal of diffuse or dense-core plaques in patients after Aducanumab treatment is not yet known, however [18]F-florbetapir PET data suggest a strong removal of amyloid (59-71%) after chronic treatment (78 weeks) in patients ^4^. These results indicate however, that treatment could be more effective at preventing plaque formation at early disease stages where amyloid is less densely aggregated.

In patients, anti-Aβ antibody treatment is associated with an increased risk of oedema (ARIA-E) and haemorrhages (ARIA-H) that could potentially be lethal in very rare cases. Several mechanisms for these side effects have been postulated, including the shifting of Aβ from the parenchyma to the blood vessels and the immune response of microglia or perivascular macrophages ^11,12,86^. The APP-SAA KI mouse model used in this study displays Prussian blue deposits with antibody treatment, but also in an age-dependent manner in untreated mice. Whether this phenotype could be related to the hTfR KI ^36^ in these mice is unclear, however another study using 5xFAD; hTfR KI mice did not observe any spontaneous microbleeds without anti-Aβ antibody treatment ^87^. We found no significant increase in Prussian Blue deposits with anti-Aβ over isotype antibody treatment.

Proteomic analysis of terminally collected CSF indicated a marked reduction in total CSF CamKIIα, Tau and α-Syn almost to baseline (untreated 3 months of age) levels, despite the remaining presence of some amyloid plaques and Lamp1^+^ neuritic dystrophy. In clinical trials, CSF total Tau levels were reduced after treatment with Aducanumab and other Aβ-targeting antibodies ^4–6^. No data on CSF total α-Syn levels after anti-Aβ antibody treatment have been reported, but increased CSF total α-Syn was previously found to be associated with increased risk of cognitive decline in non-demented adults ^88^ and reported to be increased in patients diagnosed with probable AD ^89^. Unlike patients, mouse models of amyloidosis do not develop Tau tangles, nor do they show co-pathologies such as Lewy bodies. It is therefore likely that increased CSF α-Syn, Tau and CamKIIa upon aging in APP-SAA mice reflects neuritic degeneration, rather than co-pathology ^90^. However, increased total CSF Tau and Tau phosphorylation (Ser202 and Thr205) close to plaques has been observed in the APP-SAA mouse model ^35^. Interestingly, a recent study of Aducanumab treatment in APPPS1/Tau22 mice reported less Tau phosphorylation and axonal swellings around plaques ^33^, hinting at reduced Aβ neurotoxicity in the plaque penumbra. In addition, an earlier study using laser dissection and LC-MS/MS proteomics in APPPS1 mice after Aducanumab treatment found a trend towards higher levels of Tau and α-Syn in the plaque surrounding, although these findings did not pass significance ^32^. In line with these findings, we observe a decrease in Lamp1 neurite dystrophy around residual plaques with 10 mg/kg anti-Aβ treatment. It is possible that increased microglial clustering around plaques upon anti-Aβ treatment, which is consistently reported in previous as well as the current study, could play a role by forming a protective barrier against the release of synaptotoxic Aβ fibrils or oligomers, but this hypothesis warrants further investigation ^91,92^.

CSF proteomics also indicated microglia changes upon anti-Aβ treatment, showing a dose-dependent reduction in levels of proteins such as Cst7, Cd84 and Ctsz. To investigate the effect of anti-Aβ treatment on microglial states and phenotypes in more detail, we performed bulk RNA-sequencing of isolated Cd11b^+^ microglia. We identified a relatively small, but dose-dependent effect on gene expression after chronic anti-Aβ treatment that largely showed an attenuation of the age- and pathology-associated gene expression changes in the APP-SAA KI model. Gene expression related to microglial DAM state was reduced and homeostatic gene expression was increased. Furthermore, gene set enrichment analysis (GSEA) indicated an attenuation of gene expression associated with metabolic pathways such as cholesterol homeostasis and glycolysis.

In agreement with these findings, LC-MS analysis of microglial lipid composition showed a mild reversal of the age-induced lipid changes in APP-SAA KI mice. Ganglioside GM3 levels displayed a dose-dependent reduction upon anti-Aβ treatment. GM3 was previously found to be induced with amyloid pathology in APP-SAA KI mice and is enriched in microglia that have phagocytosed MX-04^+^ Aβ ^35^. In addition, lowering of GM3 was previously found to be associated with an attenuation of amyloid pathology ^93^. In contrast, levels of cholesterol ester (CE) were increased with aging and were previously shown to be lower in MX-04^+^ vs. MX-04^-^ microglia ^35^, suggesting that CE accumulation is driven by aging rather than Aβ pathology. In line with this, CE levels were not reduced upon anti-Aβ treatment. Overall, lipidome analysis suggests that microglia do not show increased lipid burden upon chronic anti-Aβ treatment, but rather an attenuation towards pre-treatment baseline.

A reduction in microglial glycolysis was confirmed by FDG-PET, showing a dose-dependent decrease in microglial glucose uptake. In addition, a reduction in brain Trem2 and sTrem2 protein levels strongly correlated with insoluble Aβ, suggesting that the observed reduction in DAM activation is mainly associated with reduced plaque load after chronic treatment. Interestingly, previous studies that reported reduced microglial DAM activation and/or increased homeostatic gene expression after anti-Aβ treatment in mice after various dosing regimens, did so despite observing no to very little removal of Aβ ^29,30,34^. It is possible that changes in Aβ levels may have been underestimated in these studies due to the limitations of immunofluorescence analysis alone. Alternatively, Fc-gamma receptor (FcγR) stimulation may directly influence microglial DAM activation independently of Aβ, though this remains to be investigated.

Although gene expression changes overall showed an attenuation towards pre-treatment baseline, we found that anti-Aβ treatment did not lower expression of DAM genes *Lgals3* and *Spp1.* Moreover, a specific set of genes were further increased by anti-Aβ treatment in a dose-dependent manner. These genes were associated with antigen presentation (*Cd40*), cell adhesion (*Madcam1*, *Vcam1*), ROS production (*Cybb*) and cytokine signaling (*Il2rg*, *Il1rn*), with a similar trend for genes encoding chemokines such as *Cxcl10*/IP-10 and MHC-II complex proteins. *Madcam1* and *Vcam1* are classically known to encode adhesion molecules expressed in endothelial cells, but it has been reported that *Vcam1* can also be expressed by microglia ^94^. Interestingly, Il-33-induced *Vcam1* and *MHC-II* expression in microglia was previously shown to play a crucial role in chemotaxis towards plaques and subsequently promote plaque removal ^95^, suggesting this response is protective. Moreover, *Cybb* and MHC-II expression were previously identified as markers of microglia with low lipid droplet burden ^96^, indicating that extensive amyloid clearance by microglia upon anti-Aβ treatment is likely not associated with lipid droplet accumulation. Another interesting hit is the increase in *Il1rn,* encoding interleukin 1 receptor antagonist (IL1RA), which is a negative regulator of innate immune cell activation by blocking IL1 receptor activation via IL1α and IL1β ^97,98^. IL1RA has previously been shown to be associated with inhibition of NLRP3 inflammasome activation and reducing the vascular inflammatory effects of IL1β ^99^. Furthermore, *Il1rn* can be induced downstream of FcγR and SYK stimulation ^100^. Interestingly, increased IL1RN levels have also been found after TREM2 agonistic antibody treatment in mice and patients ^101,102^, suggesting this phenotype might be more broadly induced by antibody treatment that induces SYK signaling in microglia. It is likely that the increased expression of *Il1rn* is a compensatory mechanism that counteracts chronic inflammation and T-cell infiltration, but its exact function in the context of antibody treatment remains to be determined.

Finally, to validate our sequencing results and understand how these changes in microglial gene expression relate to the presence of residual plaques after chronic anti-Aβ treatment, we performed immunofluorescent staining and found an increase of markers Galectin-3 and MHC-II specifically around plaques, whereas Cd68 and Trem2 were induced around plaques similarly across treatment groups. Galectin-3 is classically known to be recruited to damaged lysosomes to promote repair ^103^. Taken together these results suggest that microglia that cluster around residual plaques maintain their DAM signature, but also acquire an antigen presentation phenotype in a dose-dependent manner, which might be associated with treatment efficiency.

Passive immunotherapy approaches have managed to bypass many of the side-effects of active immunization, but little is known about the long-term effects of this treatment on microglia, despite being key players in the efficient removal of Aβ. As anti-Aβ-antibodies are dosed chronically to an increasing number of patients, it is essential to better understand the effect of such treatments on the immune system. We find that overall microglial activation is reduced in correlation to plaque load, but microglia at residual plaques acquire a unique phenotype of both DAM and select genes associated with antigen presentation. The finding that Trem2 is still induced around residual plaques after chronic treatment indicates that microglia are still capable of actively dealing with Aβ and suggests that continued dosing with an anti-Aβ antibody could be a valid long-term treatment option, provided that treatment is given at an early time point in amyloid disease stage. Moreover, potential (co-) treatment with an agonistic Trem2 antibody might also be most effective during the earliest disease stages ^23^.

## Supplemental figures

**Figure S1:**
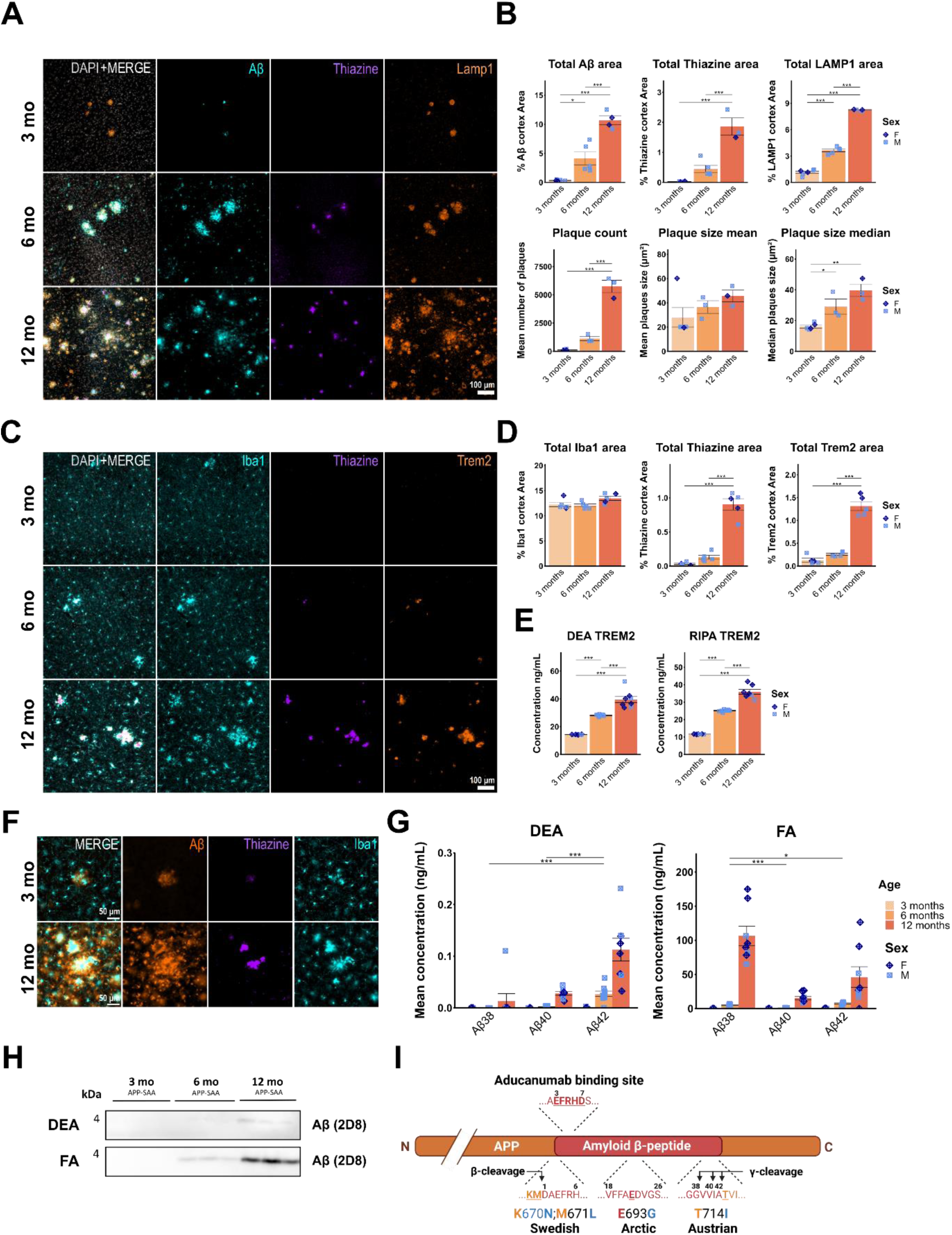
Progression of amyloid pathology in the APP-SAA mouse model. **(A)** Representative immunofluorescent images of sagittal cortical sections showing Aβ (cyan), thiazine (purple) and Lamp1 (orange) and (**B**) quantification of % cortical thiazine area, plaque count and size and % cortical Lamp1 area. (**C**) Representative immunofluorescent images of sagittal cortical sections showing Iba1 (cyan), thiazine (purple) and Trem2 (orange) and (**D**) quantification of % cortical area of Iba1, thiazine and Trem2. (**E**) ELISA quantification of Trem2 in DEA and RIPA brain lysate. (**F**) Representative immunofluorescent images showing Aβ (orange), thiazine (purple) and Iba1 (cyan), and microglia clustering around loose fibrillar Aβ plaques at 3 months and a comparison to dense plaques at 12 months. (**G**) Aβ-triplex ELISA quantification of DEA and FA extracted Aβ. (**H**) Western blot of DEA and FA fractions for Aβ (antibody 2D8). (**G**) Schematic of the human APP protein with Swedish, Artic and Austrian mutations and anti-Aβ (Aducanumab) binding site indicated. *: P < 0.05; **: P < 0.01; ***: P < 0.001. One-way ANOVA with Tukey’s post hoc test (B, D). Two-way ANOVA with Tukey’s post hoc test (G). Schematic (F) was created with BioRender.com.

**Figure S2:**
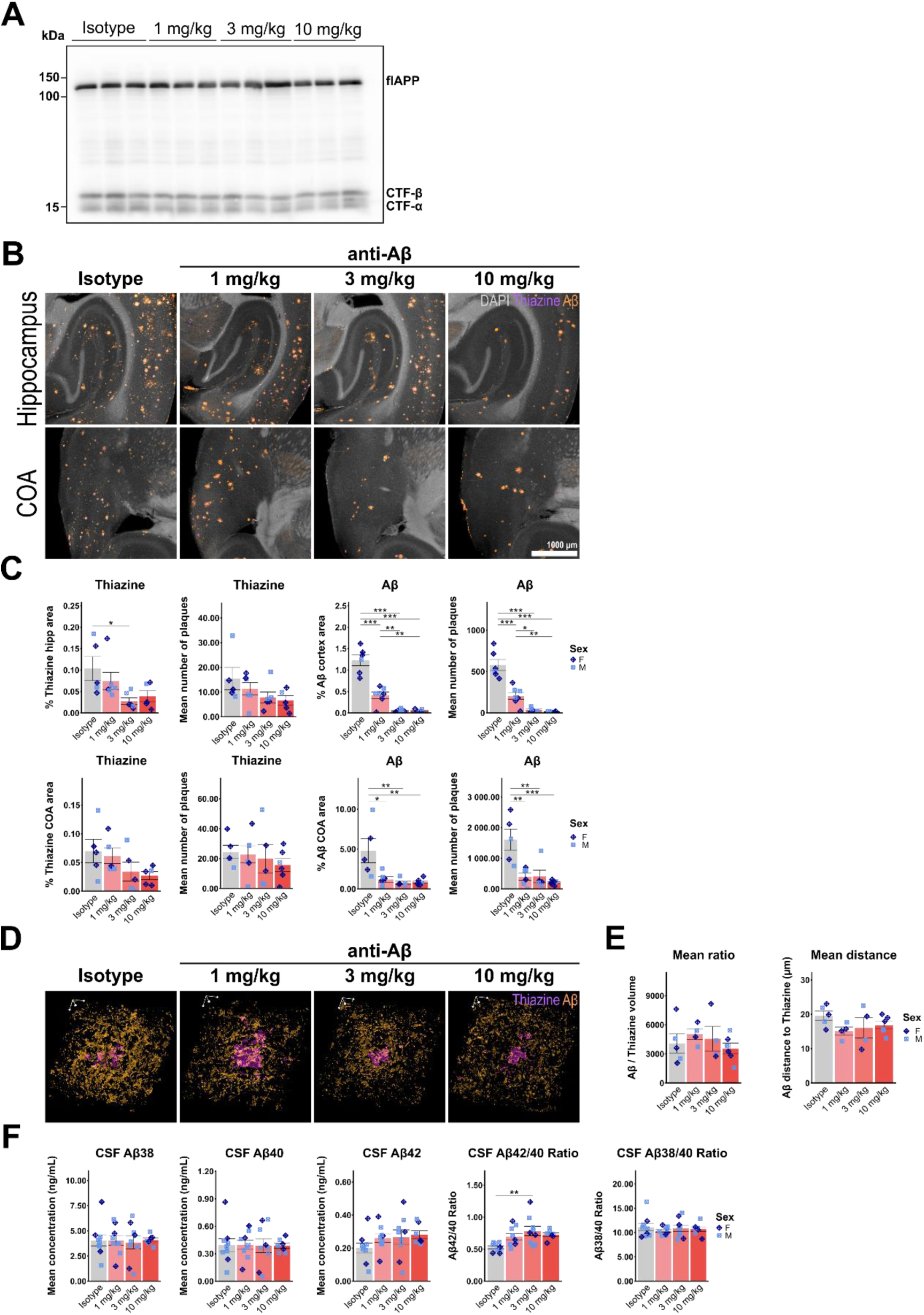
Chronic anti-Aβ treatment reduces amyloid-β levels in a dose-dependent manner. **(A)** Western blot showing full length APP, CTF-β and CTF-α in RIPA brain lysate. (**B**) Representative immunofluorescent images of sagittal hippocampus and cortico-amygdala area (COA) sections showing DAPI (grey), thiazine (purple) and Aβ (3552 antibody, orange). (**C**) Quantification of % cortical plaque and Aβ area and number in the hippocampus and COA. (**D**) Isotropic 3D rendering of confocal immunofluorescent images showing thiazine (purple) and Aβ (3552 antibody, orange). Scale bar = 10 µm. (**E**) Quantification of mean Aβ/thiazine ratio and mean distance of Aβ to thiazine border. (**F**) ELISA quantification of Aβ_38_, Aβ_40_ and Aβ_42_, as well as the Aβ_42/40 and_ Aβ_38/40_ ratio in terminally collected CSF. *: P < 0.05; **: P < 0.01; ***: P < 0.001. One-way ANOVA with Tukey’s post hoc test (C, E, F).

**Figure S3:**
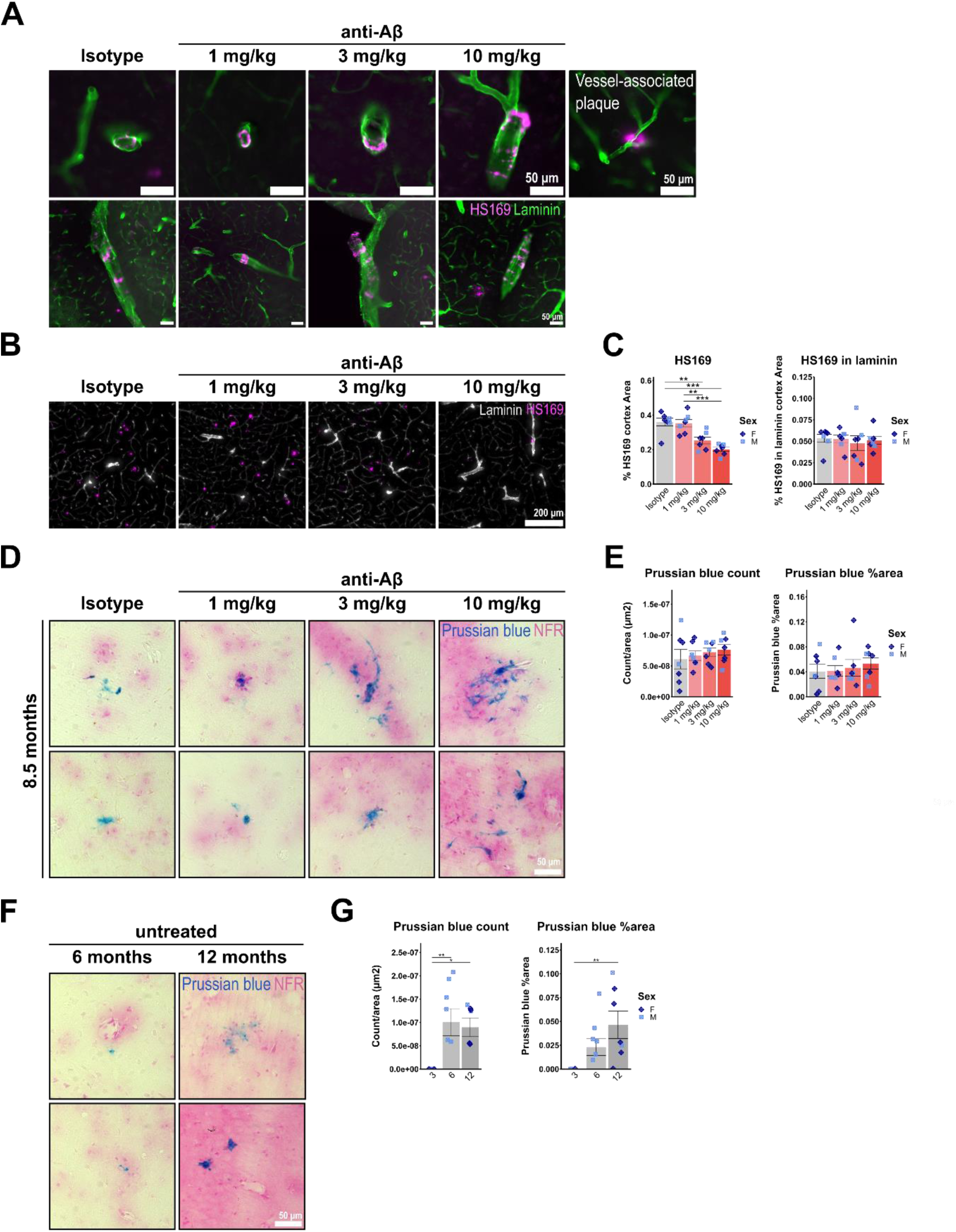
APP-SAA KI mice develop CAA and spontaneous microhaemorrhages, which are not increased by chronic anti-Aβ treatment. **(A)** Representative immunofluorescence images of CAA as stained with HS169 in APP-SAA KI mice. (**B**) Representative immunofluorescent images of sagittal cortical sections showing co-staining of Laminin and HS169. (**C**) Quantification of % cortical HS169 and HS169 in Laminin area. (**D**) Representative brightfield images of Prussian blue positive deposits and microglial-cell shaped foci in anti-Aβ or isotype treated mice. (**E**) Quantification of mean Prussian blue foci count and % area. (**F**) Representative images of Prussian blue positive deposits and microglial-cell shaped foci in 6- and 12-month-old APP-SAA KI mice. (**G**) Quantification of mean Prussian blue foci count and % area. *: P < 0.05; **: P < 0.01. One-way ANOVA with Tukey’s post hoc test (C, E, G).

**Figure S4:**
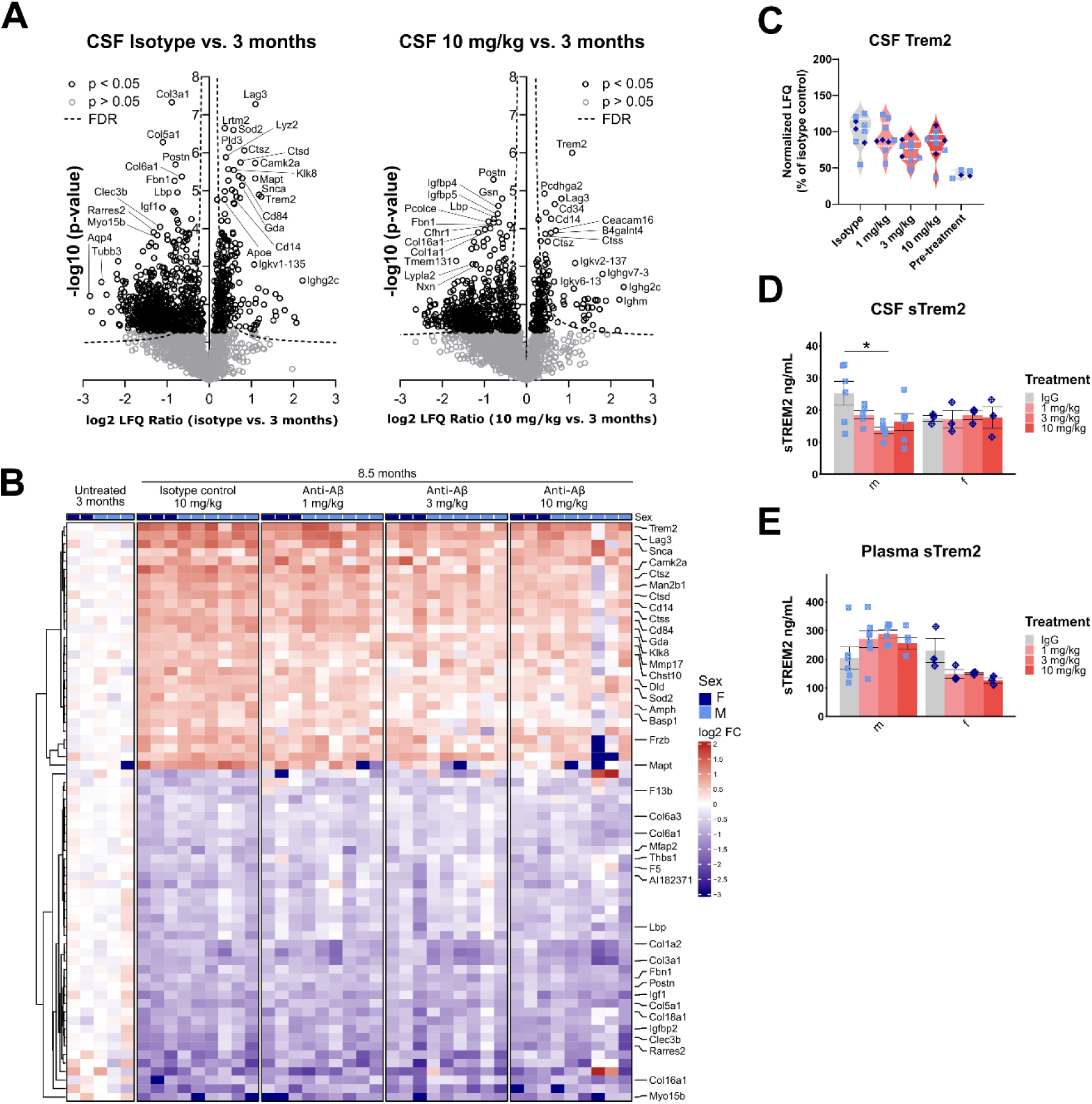
CSF proteome changes relative to pre-disease state. **(A)** Volcano plots showing upregulated and downregulated proteins in CSF comparing isotype or 10 mg/kg anti-Aβ treated animals vs. 3-month-old untreated controls. (**B**) Heatmaps showing significantly up-or downregulated proteins in isotype treated 8.5-month-old mice relative to 3-month-old untreated mice (cut-off p<0.05, log2FC < -0.5 or >0.5). (**C**) Normalized LFQ plot of Trem2 detected by LC/MS. (**D**) ELISA quantification of CSF Trem2. (**E**) ELISA quantification of terminally collected plasma Trem2. *: P < 0.05. Two-way ANOVA with Tukey’s post hoc test (D, E).

**Figure S5:**
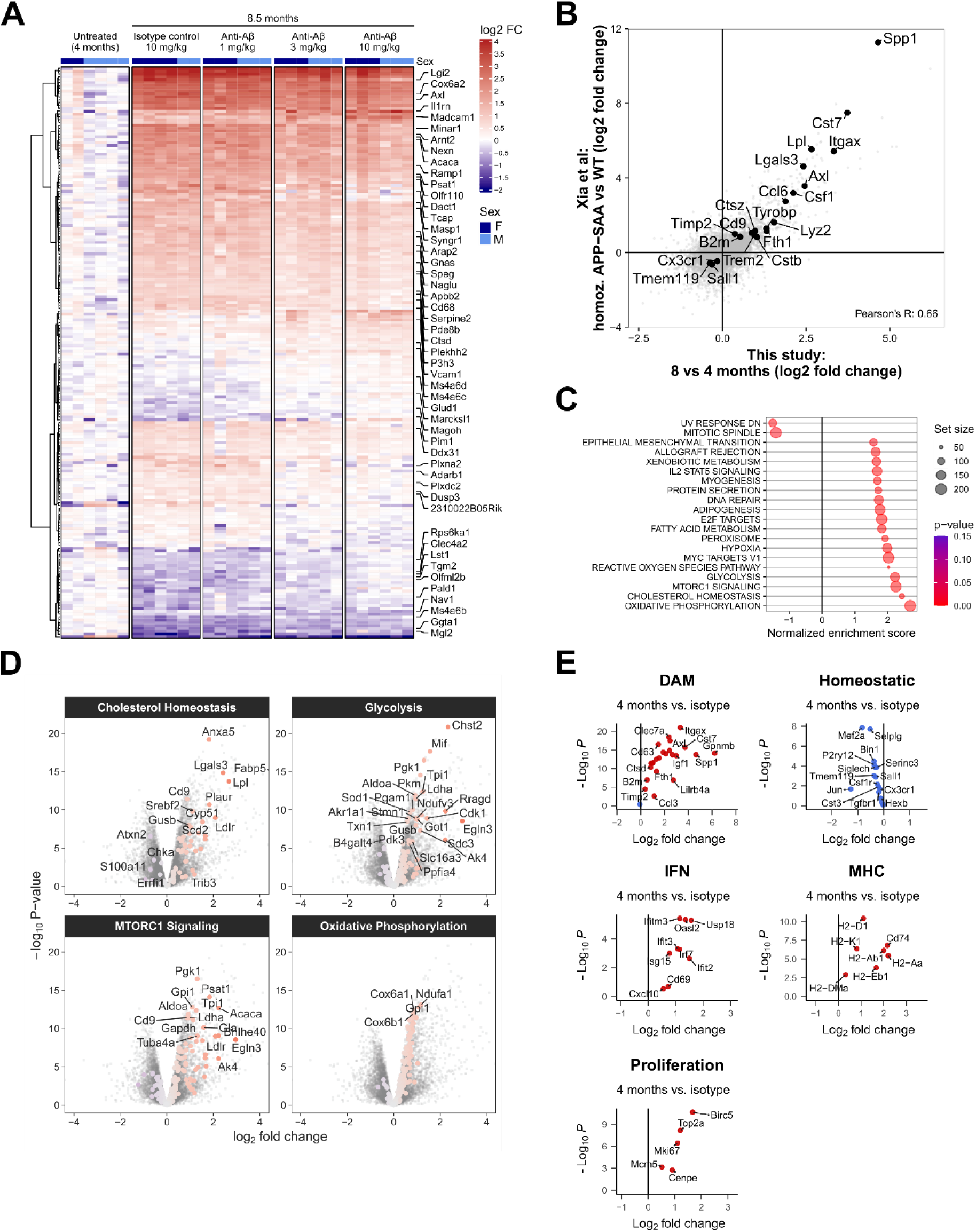
Microglial RNA-seq gene expression changes relative to pre-disease state. **(A)** Heatmap showing differentially expressed genes relative to 4-month-old untreated mice. (**B**) Correlation plot of p-values comparing gene expression changes in isolated microglia between WT vs. 8-month-old APP-SAA KI mice from Xia et. Al 2022 and changes in this study between 4-month-old untreated vs. 8-month-old isotype treated APP-SAA KI mice. (**C**) Gene set enrichment analysis (GSEA). (**D**) Volcano plots showing differentially expressed genes related to GSEA pathways. (**E**) Volcano plots showing differentially expressed genes related to microglial states from Chen and Colonna. 2021.

**Figure S6:**
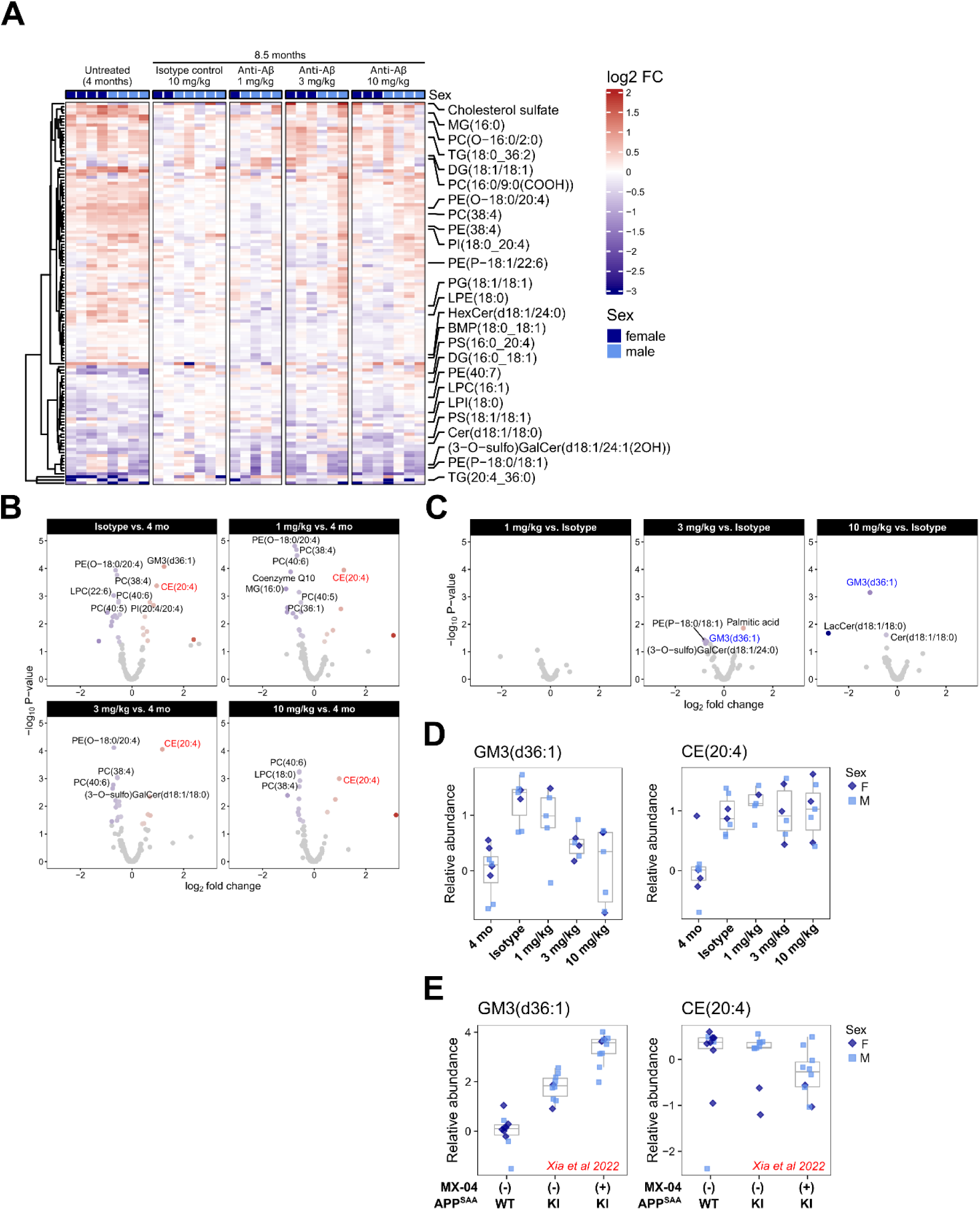
Chronic anti-Aβ treatment does not induce lipid changes in microglia. **(A)** Heatmap showing differentially regulated lipids relative to isotype control treated mice. (**B**) Volcano plots showing lipid changes in isolated microglia comparing 8-month-old treated mice vs. 4-month-old APP-SAA KI mice (**C**) Volcano plots showing lipid changes in isolated microglia comparing 8-month-old treated mice vs. isotype treated APP-SAA KI mice (**D**) Boxplots showing relative abundance of selected lipids of lipids (GM3 and CE). (**E**) Boxplots showing relative abundance of GM3 and CE from the dataset of Xia *et al.* 2022, who analysed sorted microglia from 8-month-old WT and APP-SAA mice sorted by MX-04^-^ or MX-04^+^.

**Figure S7:**
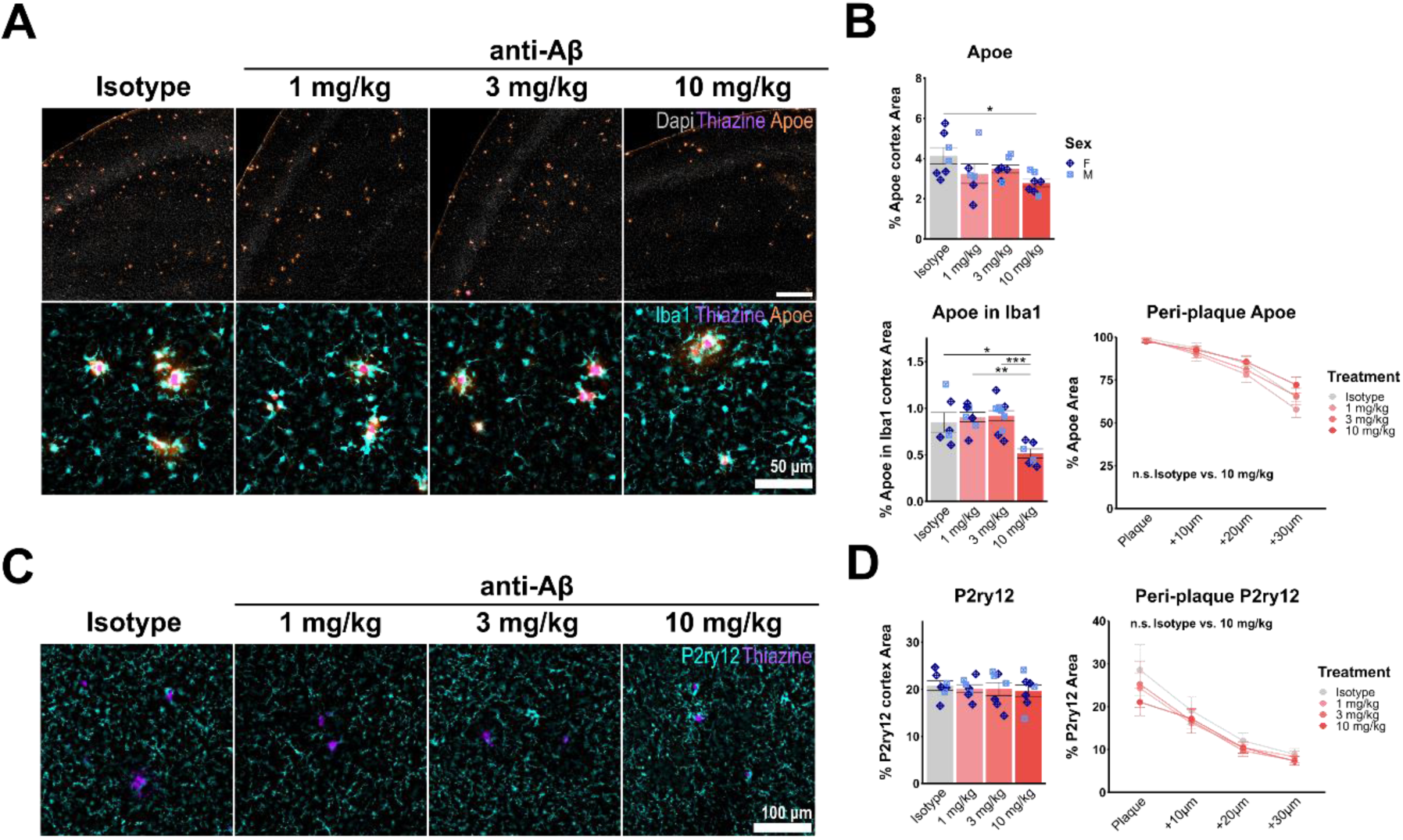
Chronic anti-Aβ treatment decreases microglial DAM activation. **(A)** Representative epifluorescence images of Dapi (grey), Iba1 (cyan), thiazine (purple) and ApoE (orange). (**B**) Quantification of % cortical ApoE and ApoE in Iba1, as well as concentric plaque analysis of ApoE in Iba1. (**C**) Representative epifluorescence images of P2ry12 (cyan) and thiazine (purple) (**D**) Quantification of % cortical P2ry12, as well as concentric plaque analysis of P2ry12. *: P < 0.05, **: P < 0.01, ***: P < 0.001. One-way ANOVA with Tukey’s post hoc test (B, D).

**Figure S8:**
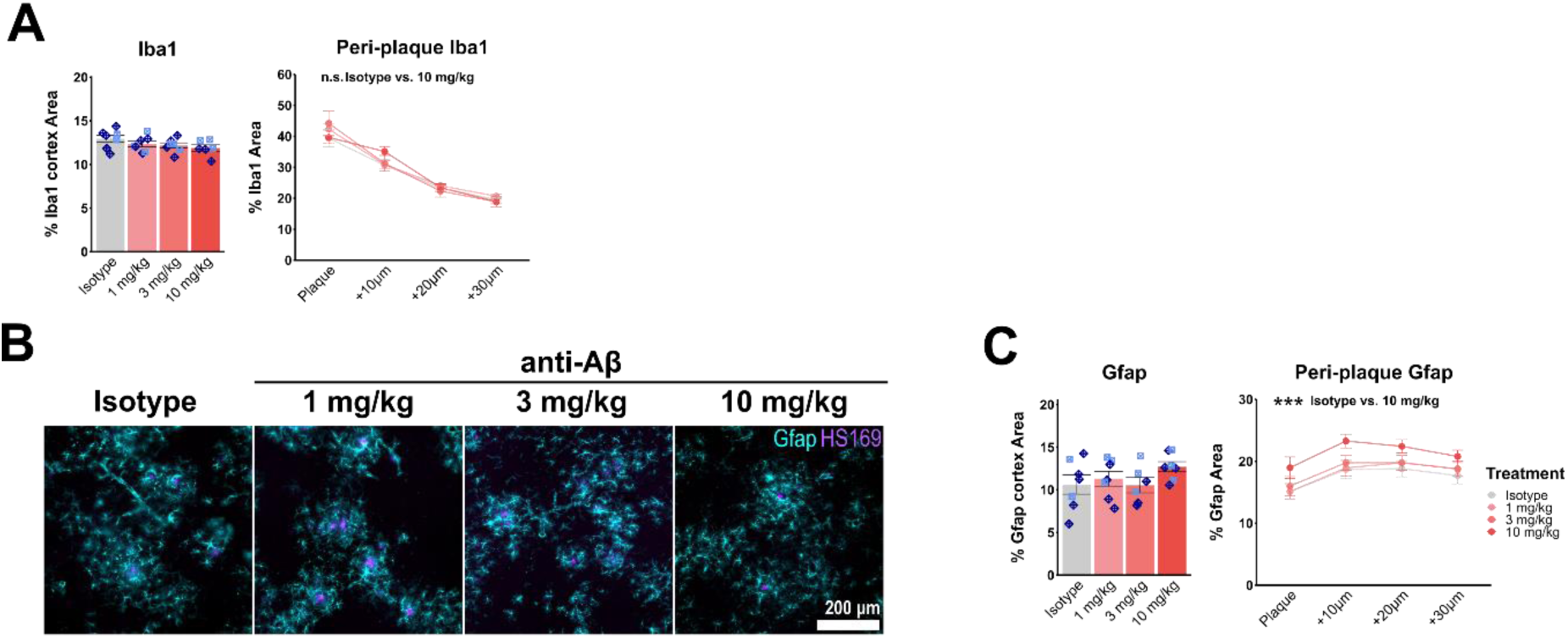
Chronic anti-Aβ treatment does not affect microgliosis and astrogliosis, but increased Gfap around residual plaques. **(A)** Quantification of % cortical Iba1 and concentric-plaque analysis of Iba1. (**B**) Representative epifluorescence images of Gfap (cyan) and thiazine (purple). (**C**) Quantification of % cortical Gfap, as well as concentric plaque analysis of Gfap. ***: P < 0.001. One-way ANOVA with Tukey’s post hoc test (A, C).

## Acknowledgements

The authors would like to thank Anne von Thaden and Manuela Schneider for their help with mouse related work. We thank Peter Nilsson for providing the HS169 compound. We thank Georg Jocher for quality control of CSF samples before LC-MS/MS analysis.

## Author contributions

L.dW., C.H., K.M.M. designed the study. M.B. designed the PET experiments, S:F:L: designed the proteomic studies. L.dW., A.F., K.S. and M.R. performed mouse experiments and collected tissue samples. S.H., R.G., S.W., A.E. (PET); S.A.M, A.B. (CSF LC-MS/MS) I.P. (image analysis); T.S., V.W., C.Ha., D.X. (RNA isolation and sequencing), S.S.D. (lipidomics), L.dW., M.E., B.N. (ELISA) and L.dW., M.E. (immunofluorescence) performed analyses. L.dW., C.H., M.B., S.F.L., J.W.L and K.M.M supervised experiments. M.W. handled legal aspects associated with animal experiments. L.dW and C.H. wrote the manuscript with input from all co-authors.

## Data availability

LC-MS/MS proteomics data have been deposited to the ProteomeXchange Consortium via the PRIDE partner repository ^104^ with the dataset identifier PXD061932. Bulk RNA-seq data have been deposited to the Gene Expression Omnibus (GEO) repository with accension number GSE288801. Data analysis scripts for reproducible analysis of bulk RNA-seq analysis have been deposited on Zenodo under digital object identifier (doi):10.5281/zenodo.14812455.

## Ethics approval

All experiments and handling of mice was performed in compliance with the German animal welfare law and with approval from the Government of Upper Bavaria (animal license: ROB-55.2-2532.Vet_02-18-32).

## Abbreviations

α-Syn: α-Synuclein
Aβ: Amyloid β-peptide
AD: Alzheimer’s disease
APP: Amyloid precursor protein
ARIA: Amyloid-related imaging abnormalities
ARIA-E: ARIA-related oedema
ARIA-H: ARIA-related haemorrhage
BCA: Bicinchoninic acid
BSA: Bovine serum albumin
CAA: Cerebral amyloid-angiopathy
CE: Cholesterol ester
COA: Cortico-amygdala area
CSF: Cerebrospinal fluid
CTF: C-terminal fragment
DAM: Disease associated microglia
DAPI: 40,6-diamidino-2-phenylindole
DEA: Diethylamine
DEG: Differentially expressed gene
diaPASEF: Data Independent Acquisition Parallel Accumulation–Serial Fragmentation
ELISA: Enzyme-linked immunosorbent assay
EtOH: Ethanol
FA: Formic acid
FcγR: Fc gamma receptor
FBB: Florbetaben
FDG: Fluorodeoxyglucose
FDR: False discovery rate
Gfap: Glial fibrillary acidic protein
GM3: Ganglioside mannose 3
GSEA: Gene set enrichment analysis
HBSS: Hanks’ buffered salt solution
hTfR: Human transferrin receptor
IFN: Intereferon
Il1rn: Interleukin-1 receptor anatagonist
i.p.: intraperitoneal
KI: Knock-in
LAMP1: Lysosomal-associated membrane protein 1
LC-MS: Liquid chromatography - mass spectrometry
LC-MS/MS: Liquid chromatography - tandem mass spectrometry
LOAD: Late-onset Alzheimer’s disease
MACS: Magnetic-activated cell sorting
MBq: Megabecquerel
MCI: Mild cognitive impairment
MHC: Major histocompatibility complex
MMF: Medetomidine-midazolam-fentanyl
MR: Magnetic resonance
MRI: Magnetic resonance imaging
MSD: Meso Scale Discovery
MX-04: Methoxy-04
NaCl: Sodium chloride
NDS: Normal donkey serum
PBS: Phosphate-buffered saline
PET: Positron-emission tomography
PFA: Paraformaldehyde
RIPA: Radioimmunoprecipitation assay
RNA-seq: RNA-sequencing
ROI: Region of interest
ROS: Reactive oxygen species
RT: Room temperature
SEM: Standard error of the mean
SUV: Standard uptake value
TBS: Tris-buffered saline
TIMS: Trapped Ion Mobility Spectrometry
Trem2: Triggering receptor expressed on myeloid cells 2
VOI: Voxel of interest
VT: Total volume of distribution

## References

1 Haass, C. & Selkoe, D. If amyloid drives Alzheimer disease, why have anti-amyloid therapies not yet slowed cognitive decline? PLoS Biol 20, e3001694, doi:10.1371/journal.pbio.3001694 (2022).

2 Selkoe, D. J. & Hardy, J. The amyloid hypothesis of Alzheimer’s disease at 25 years. EMBO Mol Med 8, 595–608, doi:10.15252/emmm.201606210 (2016).

3 Schenk, D. et al. Immunization with amyloid-beta attenuates Alzheimer-disease-like pathology in the PDAPP mouse. Nature 400, 173–177, doi:10.1038/22124 (1999).

4 Budd Haeberlein, S., et al. Two Randomized Phase 3 Studies of Aducanumab in Early Alzheimer’s Disease. The Journal of Prevention of Alzheimer’s Disease, doi:10.14283/jpad.2022.30 (2022).

5 Sims, J. R. et al. Donanemab in Early Symptomatic Alzheimer Disease: The TRAILBLAZER-ALZ 2 Randomized Clinical Trial. JAMA 330, 512–527, doi:10.1001/jama.2023.13239 (2023).

6 van Dyck, C. H. et al. Lecanemab in Early Alzheimer’s Disease. N Engl J Med 388, 9–21, doi:10.1056/NEJMoa2212948 (2023).

7 Wilcock, D. M. et al. Microglial activation facilitates Abeta plaque removal following intracranial anti-Abeta antibody administration. Neurobiol Dis 15, 11–20, doi:10.1016/j.nbd.2003.09.015 (2004).

8 Wilcock, D. M. et al. Passive amyloid immunotherapy clears amyloid and transiently activates microglia in a transgenic mouse model of amyloid deposition. J Neurosci 24, 6144–6151, doi:10.1523/JNEUROSCI.1090-04.2004 (2004).

9 Gilman, S. et al. Clinical effects of Abeta immunization (AN1792) in patients with AD in an interrupted trial. Neurology 64, 1553–1562, doi:10.1212/01.WNL.0000159740.16984.3C (2005).

10 Barakos, J. et al. Detection and Management of Amyloid-Related Imaging Abnormalities in Patients with Alzheimer’s Disease Treated with Anti-Amyloid Beta Therapy. J Prev Alzheimers Dis 9, 211–220, doi:10.14283/jpad.2022.21 (2022).

11 Taylor, X. et al. Amyloid-beta (Abeta) immunotherapy induced microhemorrhages are linked to vascular inflammation and cerebrovascular damage in a mouse model of Alzheimer’s disease. Mol Neurodegener 19, 77, doi:10.1186/s13024-024-00758-0 (2024).

12 Taylor, X. et al. Amyloid-beta (Abeta) immunotherapy induced microhemorrhages are associated with activated perivascular macrophages and peripheral monocyte recruitment in Alzheimer’s disease mice. Mol Neurodegener 18, 59, doi:10.1186/s13024-023-00649-w (2023).

13 Schlepckow, K., Morenas-Rodriguez, E., Hong, S. & Haass, C. Stimulation of TREM2 with agonistic antibodies-an emerging therapeutic option for Alzheimer’s disease. Lancet Neurol 22, 1048–1060, doi:10.1016/S1474-4422(23)00247-8 (2023).

14 Lewcock, J. W. et al. Emerging Microglia Biology Defines Novel Therapeutic Approaches for Alzheimer’s Disease. Neuron 108, 801–821, doi:10.1016/j.neuron.2020.09.029 (2020).

15 Guerreiro, R. et al. TREM2 variants in Alzheimer’s disease. N Engl J Med 368, 117–127, doi:10.1056/NEJMoa1211851 (2013).

16 Gratuze, M., Leyns, C. E. G. & Holtzman, D. M. New insights into the role of TREM2 in Alzheimer’s disease. Mol Neurodegener 13, 66, doi:10.1186/s13024-018-0298-9 (2018).

17 Krasemann, S. et al. The TREM2-APOE Pathway Drives the Transcriptional Phenotype of Dysfunctional Microglia in Neurodegenerative Diseases. Immunity 47, 566–581 e569, doi:10.1016/j.immuni.2017.08.008 (2017).

18 Keren-Shaul, H. et al. A Unique Microglia Type Associated with Restricting Development of Alzheimer’s Disease. Cell 169, 1276–1290 e1217, doi:10.1016/j.cell.2017.05.018 (2017).

19 Zhao, Y. et al. TREM2 Is a Receptor for beta-Amyloid that Mediates Microglial Function. Neuron 97, 1023–1031 e1027, doi:10.1016/j.neuron.2018.01.031 (2018).

20 Parhizkar, S. et al. Loss of TREM2 function increases amyloid seeding but reduces plaque-associated ApoE. Nat Neurosci 22, 191–204, doi:10.1038/s41593-018-0296-9 (2019).

21 Mazaheri, F. et al. TREM2 deficiency impairs chemotaxis and microglial responses to neuronal injury. EMBO Rep 18, 1186–1198, doi:10.15252/embr.201743922 (2017).

22 Jonsson, T. et al. Variant of TREM2 associated with the risk of Alzheimer’s disease. N Engl J Med 368, 107–116, doi:10.1056/NEJMoa1211103 (2013).

23 Morenas-Rodríguez, E. et al. Soluble TREM2 in CSF and its association with other biomarkers and cognition in autosomal-dominant Alzheimer’s disease: a longitudinal observational study. The Lancet Neurology 21, 329–341, doi:10.1016/s1474-4422(22)00027-8 (2022).

24 Ewers, M. et al. Higher CSF sTREM2 and microglia activation are associated with slower rates of beta-amyloid accumulation. EMBO Mol Med 12, e12308, doi:10.15252/emmm.202012308 (2020).

25 Ewers, M. et al. Increased soluble TREM2 in cerebrospinal fluid is associated with reduced cognitive and clinical decline in Alzheimer’s disease. Sci Transl Med 11, doi:10.1126/scitranslmed.aav6221 (2019).

26 Franzmeier, N. et al. Higher CSF sTREM2 attenuates ApoE4-related risk for cognitive decline and neurodegeneration. Molecular Neurodegeneration 15, doi:10.1186/s13024-020-00407-2 (2020).

27 Xiang, X. et al. TREM2 deficiency reduces the efficacy of immunotherapeutic amyloid clearance. EMBO Mol Med 8, 992–1004, doi:10.15252/emmm.201606370 (2016).

28 Wilcock, D. M. et al. Diverse inflammatory responses in transgenic mouse models of Alzheimer’s disease and the effect of immunotherapy on these responses. ASN Neuro 3, 249–258, doi:10.1042/AN20110018 (2011).

29 Cadiz, M. P. et al. Aducanumab anti-amyloid immunotherapy induces sustained microglial and immune alterations. J Exp Med 221, doi:10.1084/jem.20231363 (2024).

30 Xiong, M. et al. APOE immunotherapy reduces cerebral amyloid angiopathy and amyloid plaques while improving cerebrovascular function. Sci Transl Med 13, doi:10.1126/scitranslmed.abd7522 (2021).

31 Sevigny, J. et al. The antibody aducanumab reduces Abeta plaques in Alzheimer’s disease. Nature 537, 50–56, doi:10.1038/nature19323 (2016).

32 Bastrup, J. et al. Anti-Abeta Antibody Aducanumab Regulates the Proteome of Senile Plaques and Closely Surrounding Tissue in a Transgenic Mouse Model of Alzheimer’s Disease. J Alzheimers Dis 79, 249–265, doi:10.3233/JAD-200715 (2021).

33 Welikovitch, L. A. et al. Tau, synapse loss and gliosis progress in an Alzheimer’s mouse model after amyloid-beta immunotherapy. Brain, doi:10.1093/brain/awae345 (2024).

34 Da Mesquita, S. et al. Meningeal lymphatics affect microglia responses and anti-Abeta immunotherapy. Nature, doi:10.1038/s41586-021-03489-0 (2021).

35 Xia, D. et al. Novel App knock-in mouse model shows key features of amyloid pathology and reveals profound metabolic dysregulation of microglia. Mol Neurodegener 17, 41, doi:10.1186/s13024-022-00547-7 (2022).

36 Kariolis, M. S. et al. Brain delivery of therapeutic proteins using an Fc fragment blood-brain barrier transport vehicle in mice and monkeys. Sci Transl Med 12, doi:10.1126/scitranslmed.aay1359 (2020).

37 van Lengerich, B. et al. A TREM2-activating antibody with a blood-brain barrier transport vehicle enhances microglial metabolism in Alzheimer’s disease models. Nat Neurosci, doi:10.1038/s41593-022-01240-0 (2023).

38 Hudziak, R. M. et al. p185HER2 monoclonal antibody has antiproliferative effects in vitro and sensitizes human breast tumor cells to tumor necrosis factor. Mol Cell Biol 9, 1165–1172, doi:10.1128/mcb.9.3.1165-1172.1989 (1989).

39 Overhoff, F. et al. Automated Spatial Brain Normalization and Hindbrain White Matter Reference Tissue Give Improved [(18)F]-Florbetaben PET Quantitation in Alzheimer’s Model Mice. Front Neurosci 10, 45, doi:10.3389/fnins.2016.00045 (2016).

40 Reifschneider, A. et al. Loss of TREM2 rescues hyperactivation of microglia, but not lysosomal deficits and neurotoxicity in models of progranulin deficiency. EMBO J, e109108, doi:10.15252/embj.2021109108 (2022).

41 Schiffer, W. K., Mirrione, M. M. & Dewey, S. L. Optimizing experimental protocols for quantitative behavioral imaging with 18F-FDG in rodents. J Nucl Med 48, 277–287 (2007).

42 Xiang, X. et al. Microglial activation states drive glucose uptake and FDG-PET alterations in neurodegenerative diseases. Sci Transl Med 13, eabe5640, doi:10.1126/scitranslmed.abe5640 (2021).

43 Logan, J. et al. Graphical analysis of reversible radioligand binding from time-activity measurements applied to [N-11C-methyl]-(-)-cocaine PET studies in human subjects. J Cereb Blood Flow Metab 10, 740–747, doi:10.1038/jcbfm.1990.127 (1990).

44 Ma, Y. et al. A three-dimensional digital atlas database of the adult C57BL/6J mouse brain by magnetic resonance microscopy. Neuroscience 135, 1203–1215, doi:10.1016/j.neuroscience.2005.07.014 (2005).

45 Pesamaa, I. et al. A microglial activity state biomarker panel differentiates FTD-granulin and Alzheimer’s disease patients from controls. Mol Neurodegener 18, 70, doi:10.1186/s13024-023-00657-w (2023).

46 Schindelin, J. et al. Fiji: an open-source platform for biological-image analysis. Nat Methods 9, 676–682, doi:10.1038/nmeth.2019 (2012).

47 Haase, R. et al. CLIJ: GPU-accelerated image processing for everyone. Nat Methods 17, 5–6, doi:10.1038/s41592-019-0650-1 (2020).

48 R Core Team. R: A language and environment for statistical computing. R Foundation for Statistical Computing, Vienna, Austria. URL: https://www.R-project.org/. (2023).

49 Ollion, J., Cochennec, J., Loll, F., Escude, C. & Boudier, T. TANGO: a generic tool for high-throughput 3D image analysis for studying nuclear organization. Bioinformatics 29, 1840–1841, doi:10.1093/bioinformatics/btt276 (2013).

50 napari: a multi-dimensional image viewer for python. doi:10.5281/zenodo.3555620 (2019).

51 Willem, M. et al. eta-Secretase processing of APP inhibits neuronal activity in the hippocampus. Nature 526, 443–447, doi:10.1038/nature14864 (2015).

52 Schlepckow, K. et al. Enhancing protective microglial activities with a dual function TREM2 antibody to the stalk region. EMBO Mol Med 12, e11227, doi:10.15252/emmm.201911227 (2020).

53 Hughes, C. S. et al. Single-pot, solid-phase-enhanced sample preparation for proteomics experiments. Nat Protoc 14, 68–85, doi:10.1038/s41596-018-0082-x (2019).

54 Demichev, V., Messner, C. B., Vernardis, S. I., Lilley, K. S. & Ralser, M. DIA-NN: neural networks and interference correction enable deep proteome coverage in high throughput. Nat Methods 17, 41–44, doi:10.1038/s41592-019-0638-x (2020).

55 Tyanova, S. et al. The Perseus computational platform for comprehensive analysis of (prote)omics data. Nat Methods 13, 731–740, doi:10.1038/nmeth.3901 (2016).

56 Tusher, V. G., Tibshirani, R. & Chu, G. Significance analysis of microarrays applied to the ionizing radiation response. Proceedings of the National Academy of Sciences 98, 5116–5121, doi:10.1073/pnas.091062498 (2001).

57 Ewels, P. A. et al. The nf-core framework for community-curated bioinformatics pipelines. Nat Biotechnol 38, 276–278, doi:10.1038/s41587-020-0439-x (2020).

58 Dobin, A. et al. STAR: ultrafast universal RNA-seq aligner. Bioinformatics 29, 15–21, doi:10.1093/bioinformatics/bts635 (2013).

59 Patro, R., Duggal, G., Love, M. I., Irizarry, R. A. & Kingsford, C. Salmon provides fast and bias-aware quantification of transcript expression. Nat Methods 14, 417–419, doi:10.1038/nmeth.4197 (2017).

60 Di Tommaso, P. et al. Nextflow enables reproducible computational workflows. Nat Biotechnol 35, 316–319, doi:10.1038/nbt.3820 (2017).

61 Law, C. W., Chen, Y., Shi, W. & Smyth, G. K. voom: Precision weights unlock linear model analysis tools for RNA-seq read counts. Genome Biol 15, R29, doi:10.1186/gb-2014-15-2-r29 (2014).

62 Robinson, M. D. & Oshlack, A. A scaling normalization method for differential expression analysis of RNA-seq data. Genome Biol 11, R25, doi:10.1186/gb-2010-11-3-r25 (2010).

63 Ritchie, M. E. et al. limma powers differential expression analyses for RNA-sequencing and microarray studies. Nucleic Acids Res 43, e47, doi:10.1093/nar/gkv007 (2015).

64 Benjamini, Y. & Hochberg, Y. Controlling the False Discovery Rate: A Practical and Powerful Approach to Multiple Testing. Journal of the Royal Statistical Society Series B: Statistical Methodology 57, 289–300, doi:10.1111/j.2517-6161.1995.tb02031.x (1995).

65 Korotkevich, G., et al. Fast gene set enrichment analysis. *BioRxiv*, doi:10.1101/060012 (2019).

66 Ahlmann-Eltze, C. & Patil, I. ggsignif: R Package for Displaying Significance Brackets for ’ggplot2’. doi:10.31234/osf.io/7awm6.

67 Wickham, H. et al. Welcome to the Tidyverse. Journal of Open Source Software 4, doi:10.21105/joss.01686 (2019).

68 Mena, R., Edwards, P., Perez-Olvera, O. & Wischik, C. M. Monitoring pathological assembly of tau and beta-amyloid proteins in Alzheimer’s disease. Acta Neuropathol 89, 50–56, doi:10.1007/BF00294259 (1995).

69 Biechele, G. et al. Glitter in the Darkness? Nonfibrillar beta-Amyloid Plaque Components Significantly Impact the beta-Amyloid PET Signal in Mouse Models of Alzheimer Disease. J Nucl Med 63, 117–124, doi:10.2967/jnumed.120.261858 (2022).

70 Frank, S. et al. TREM2 is upregulated in amyloid plaque-associated microglia in aged APP23 transgenic mice. Glia 56, 1438–1447, doi:10.1002/glia.20710 (2008).

71 Sebastian Monasor, L., et al. Fibrillar Abeta triggers microglial proteome alterations and dysfunction in Alzheimer mouse models. Elife 9, doi:10.7554/eLife.54083 (2020).

72 Devkota, S., Williams, T. D. & Wolfe, M. S. Familial Alzheimer’s disease mutations in amyloid protein precursor alter proteolysis by gamma-secretase to increase amyloid beta-peptides of >/=45 residues. J Biol Chem 296, 100281, doi:10.1016/j.jbc.2021.100281 (2021).

73 Reinert, J. et al. Abeta38 in the brains of patients with sporadic and familial Alzheimer’s disease and transgenic mouse models. J Alzheimers Dis 39, 871–881, doi:10.3233/JAD-131373 (2014).

74 Dimitrov, M. et al. Alzheimer’s disease mutations in APP but not gamma-secretase modulators affect epsilon-cleavage-dependent AICD production. Nat Commun 4, 2246, doi:10.1038/ncomms3246 (2013).

75 Arndt, J. W. et al. Structural and kinetic basis for the selectivity of aducanumab for aggregated forms of amyloid-β. Scientific Reports 8, doi:10.1038/s41598-018-24501-0 (2018).

76 Brendel, M. et al. Cross-sectional comparison of small animal [18F]-florbetaben amyloid-PET between transgenic AD mouse models. PLoS One 10, e0116678, doi:10.1371/journal.pone.0116678 (2015).

77 Klingstedt, T. et al. Dual-ligand fluorescence microscopy enables chronological and spatial histological assignment of distinct amyloid-beta deposits. J Biol Chem 301, 108032, doi:10.1016/j.jbc.2024.108032 (2025).

78 Hansson, O. Biomarkers for neurodegenerative diseases. Nat Med 27, 954–963, doi:10.1038/s41591-021-01382-x (2021).

79 Selkoe, D. J. The advent of Alzheimer treatments will change the trajectory of human aging. Nat Aging 4, 453–463, doi:10.1038/s43587-024-00611-5 (2024).

80 Chen, Y. & Colonna, M. Microglia in Alzheimer’s disease at single-cell level. Are there common patterns in humans and mice? J Exp Med 218, doi:10.1084/jem.20202717 (2021).

81 Ulland, T. K. et al. TREM2 Maintains Microglial Metabolic Fitness in Alzheimer’s Disease. Cell 170, 649–663 e613, doi:10.1016/j.cell.2017.07.023 (2017).

82 Feiten, A. F. et al. TREM2 expression level is critical for microglial state, metabolic capacity and efficacy of TREM2 agonism. doi:10.1101/2024.07.18.604115 (2024).

83 Plowey, E. D. et al. Alzheimer disease neuropathology in a patient previously treated with aducanumab. Acta Neuropathol 144, 143–153, doi:10.1007/s00401-022-02433-4 (2022).

84 Soderberg, L. et al. Lecanemab, Aducanumab, and Gantenerumab - Binding Profiles to Different Forms of Amyloid-Beta Might Explain Efficacy and Side Effects in Clinical Trials for Alzheimer’s Disease. Neurotherapeutics 20, 195–206, doi:10.1007/s13311-022-01308-6 (2023).

85 Hock, C. & Nitsch, R. M. Clinical observations with AN-1792 using TAPIR analyses. Neurodegener Dis 2, 273–276, doi:10.1159/000090368 (2005).

86 Wilcock, D. M. et al. Passive immunotherapy against Abeta in aged APP-transgenic mice reverses cognitive deficits and depletes parenchymal amyloid deposits in spite of increased vascular amyloid and microhemorrhage. J Neuroinflammation 1, 24, doi:10.1186/1742-2094-1-24 (2004).

87 Pizzo, M. E. et al. Engineering anti-amyloid antibodies with transferrin receptor targeting improves brain biodistribution and mitigates ARIA. doi:10.1101/2024.07.26.604664 (2024).

88 Liu, W. et al. Cerebrospinal fluid alpha-synuclein adds the risk of cognitive decline and is associated with tau pathology among non-demented older adults. Alzheimers Res Ther 16, 103, doi:10.1186/s13195-024-01463-2 (2024).

89 Majbour, N. K. et al. Increased levels of CSF total but not oligomeric or phosphorylated forms of alpha-synuclein in patients diagnosed with probable Alzheimer’s disease. Sci Rep 7, 40263, doi:10.1038/srep40263 (2017).

90 Uytterhoeven, V., Verstreken, P. & Nachman, E. Synaptic sabotage: How Tau and alpha-Synuclein undermine synaptic health. J Cell Biol 224, doi:10.1083/jcb.202409104 (2025).

91 Fruhwurth, S., Zetterberg, H. & Paludan, S. R. Microglia and amyloid plaque formation in Alzheimer’s disease - Evidence, possible mechanisms, and future challenges. J Neuroimmunol 390, 578342, doi:10.1016/j.jneuroim.2024.578342 (2024).

92 Stern, A. M. et al. Abundant Abeta fibrils in ultracentrifugal supernatants of aqueous extracts from Alzheimer’s disease brains. Neuron 111, 2012–2020 e2014, doi:10.1016/j.neuron.2023.04.007 (2023).

93 Dodge, J. C. et al. Glucosylceramide synthase inhibition reduces ganglioside GM3 accumulation, alleviates amyloid neuropathology, and stabilizes remote contextual memory in a mouse model of Alzheimer’s disease. Alzheimers Res Ther 14, 19, doi:10.1186/s13195-022-00966-0 (2022).

94 Chabot, S., Williams, G. & Yong, V. W. Microglial production of TNF-alpha is induced by activated T lymphocytes. Involvement of VLA-4 and inhibition by interferonbeta-1b. J Clin Invest 100, 604–612, doi:10.1172/JCI119571 (1997).

95 Lau, S. F. et al. The VCAM1-ApoE pathway directs microglial chemotaxis and alleviates Alzheimer’s disease pathology. Nat Aging 3, 1219–1236, doi:10.1038/s43587-023-00491-1 (2023).

96 Marschallinger, J. et al. Lipid-droplet-accumulating microglia represent a dysfunctional and proinflammatory state in the aging brain. Nat Neurosci 23, 194–208, doi:10.1038/s41593-019-0566-1 (2020).

97 Hannum, C. H. et al. Interleukin-1 receptor antagonist activity of a human interleukin-1 inhibitor. Nature 343, 336–340, doi:10.1038/343336a0 (1990).

98 Wang, Y. et al. Identification of an IL-1 receptor mutation driving autoinflammation directs IL-1-targeted drug design. Immunity 56, 1485–1501 e1487, doi:10.1016/j.immuni.2023.05.014 (2023).

99 Iannitti, R. G. et al. IL-1 receptor antagonist ameliorates inflammasome-dependent inflammation in murine and human cystic fibrosis. Nat Commun 7, 10791, doi:10.1038/ncomms10791 (2016).

100 Becker, K. L. et al. Aspergillus Cell Wall Chitin Induces Anti- and Proinflammatory Cytokines in Human PBMCs via the Fc-gamma Receptor/Syk/PI3K Pathway. mBio 7, doi:10.1128/mBio.01823-15 (2016).

101 Long, H. et al. Preclinical and first-in-human evaluation of AL002, a novel TREM2 agonistic antibody for Alzheimer’s disease. Alzheimers Res Ther 16, 235, doi:10.1186/s13195-024-01599-1 (2024).

102 Price, B. R. et al. Therapeutic Trem2 activation ameliorates amyloid-beta deposition and improves cognition in the 5XFAD model of amyloid deposition. J Neuroinflammation 17, 238, doi:10.1186/s12974-020-01915-0 (2020).

103 Jia, J. et al. Galectin-3 Coordinates a Cellular System for Lysosomal Repair and Removal. Dev Cell 52, 69–87 e68, doi:10.1016/j.devcel.2019.10.025 (2020).

104 Perez-Riverol, Y. et al. The PRIDE database at 20 years: 2025 update. Nucleic Acids Res 53, D543–D553, doi:10.1093/nar/gkae1011 (2025).

